# From Proteome Mining to Structural Validation: Phosphopyruvate Hydratase as a Structurally Tractable Drug Target in Kinetoplastid Parasites

**DOI:** 10.64898/2026.06.09.731156

**Authors:** Luis Daniel Goyzueta-Mamani, Haruna Luz Barazorda-Ccahuana, Michelle G. Ng, Laura Pineda, José L. Medina-Franco, Mónica Florin-Christensen, Eduardo Antonio Ferraz Coelho, Carmenza Spadafora, Miguel Angel Chávez-Fumagalli

## Abstract

Chagas disease, caused by *Trypanosoma cruzi*, demands novel therapeutic strategies that overcome the toxicity and limited efficacy of current treatments. To address this need, herein we report an integrative, target-centric strategy that combines parasite proteome mining, structural modeling, and experimental validation. Functional enrichment and druggability analyses identified phosphopyruvate hydratase (PPH) as a promising candidate due to its essential metabolic role and limited similarity to human homologs. Notably, proteome mining revealed the presence and conservation of PPH across kinetoplastid parasites, including *Leishmania donovani*, supporting its evaluation beyond *T. cruzi*. For the selected PPH sequences, AlphaFold-derived three-dimensional models underwent extensive molecular dynamics refinement, yielding stable conformational ensembles suitable for structure-based studies. Using this validated model, virtual screening of the Latin American Natural Products Database - LANaPDB - identified aptosimon as a top-ranked compound candidate. Molecular dynamics simulations further showed ligand-dependent binding behavior, suggesting alternative binding modes distinct from the canonical substrate configuration. *In vitro* assays demonstrated consistent antiparasitic activity against intracellular *T. cruzi* amastigotes (IC₅₀ = 3.52 ± 0.023 µg/mL) and *Leishmania donovani* promastigotes (IC₅₀ = 13.06 ± 0.018 µg/mL), supporting the biological relevance of the aptosimon-related lignan chemotype, hinokinin, across two kinetoplastid parasite models. Together, these results support PPH as a structurally tractable and biologically relevant candidate target, while identifying an aptosimon-related lignan chemotype, represented experimentally by hinokinin, as a cross-species antiparasitic scaffold that warrants further biochemical target-validation studies.

## Introduction

Chagas disease, caused by the protozoan parasite *Trypanosoma cruzi* [1], remains a major public health concern, particularly in Latin America, with an estimated 6–7 million people infected worldwide. Although traditionally confined to endemic regions, population mobility has led to an increasing number of cases in non-endemic areas, including North America and Europe [2,3]. The disease progresses through an acute phase, often asymptomatic or mildly symptomatic, followed by a chronic phase that can lead to severe cardiac and gastrointestinal complications, including cardiomyopathy, arrhythmias, and megasyndromes [4]. Despite its clinical impact, Chagas disease continues to be classified as a neglected tropical disease (NTD), disproportionately affecting populations with limited access to healthcare and early diagnosis [5,6].

Current therapeutic options for Chagas disease are limited to benznidazole and nifurtimox, drugs that were developed decades ago and present significant limitations [7]. Even though both drugs are used in clinical settings as antiparasitic medicines, their exact molecular targets and full mechanisms of action are yet unknown. Both function as prodrugs that provide cytotoxic effects when activated by *T. cruzi* nitroreductases. Their limitations include long treatment regimens, severe adverse effects, and reduced efficacy in the chronic phase of infection. Furthermore, variability in patient response and the potential emergence of drug resistance complicate disease management. The absence of an effective vaccine further underscores the urgent need for alternative therapeutic strategies [8]. In this context, the identification of novel molecular targets that enable selective parasite inhibition while minimizing host toxicity represents a critical priority in antiparasitic drug discovery.

Advances in systems biology and bioinformatics have enabled the use of proteome-wide approaches to systematically identify potential drug targets in parasitic organisms [9]. Proteome mining strategies, combined with functional enrichment and druggability assessment, allow for the prioritization of proteins that are essential for parasite survival and sufficiently divergent from human homologs [10]. This integrative approach facilitates the identification of targets that not only play central roles in parasite metabolism but also offer opportunities for selective therapeutic intervention.

Several parasite enzymes involved in sterol biosynthesis, redox metabolism, nucleic acid processing, and central carbon metabolism have been explored as drug-discovery targets in kinetoplastids. Examples include enzymes associated with ergosterol biosynthesis, trypanothione-dependent redox homeostasis, nucleic acid metabolism, and glycolysis [11]. These targets illustrate how essential parasite pathways can provide opportunities for selective therapeutic intervention when combined with host-similarity filtering and structure-based analysis. Within this broader target-discovery context, phosphopyruvate hydratase/enolase was considered as part of a proteome-guided strategy to identify metabolically relevant and structurally tractable parasite vulnerabilities [12].

In parallel, natural products have continuously shown to be a valuable source of chemically diverse and biologically active compounds for antiparasitic drug discovery [13]. Phytochemicals such as flavonoids, alkaloids, terpenoids, and bufadienolides have demonstrated trypanocidal and leishmanicidal activity [14–17], highlighting their potential as starting points for drug development [18]. Their structural diversity and evolutionary optimization enable interactions with biological targets distinct from those of conventional synthetic compounds, potentially improving selectivity and reducing the likelihood of resistance [19]. Curated natural product databases, including regionally focused repositories such as the Latin American Natural Product Database (LANaPDB) [20] and its associated collections, provide organized and chemically annotated resources that facilitate systematic virtual screening, scaffold prioritization, and integration with structure-based drug discovery workflows. Integrating natural product libraries with structure-based approaches, therefore provides a promising framework for identifying novel antiparasitic scaffolds.

In this study, we applied an integrative strategy combining proteome mining, structural modeling, and experimental validation to investigate phosphopyruvate hydratase as a structurally tractable candidate drug target in kinetoplastid parasites. Starting from proteome-wide analyses in *T. cruzi*, PPH was prioritized and modeled in three dimensions using computational approaches. The resulting structure was refined and validated through molecular dynamics (MD) simulations, enabling the identification of potential binding sites and the structure-based screening of natural product libraries. Proteome-guided analysis further revealed conservation of PPH in *L. donovani*, supporting cross-species evaluation. Selected natural product compounds were consequently assessed both computationally and experimentally *in vitro* assays to determine their antiparasitic activity and broader relevance. This combined framework provides a comprehensive evaluation of PPH as a structurally validated target and offers new insights into natural product-derived scaffolds with activity against kinetoplastid parasites.

## **3.** Materials and Methods

### 3.1 Proteome Mining and Enrichment Analysis

The reference proteome of *T. cruzi* (Proteome ID: UP000002296) was retrieved from the UniProt database [21] (http://www.uniprot.org; accessed 10 November 2024) and used as the initial dataset for target discovery. All protein entries were imported into the R statistical environment (version 4.1.0) for functional annotation, enrichment analysis, and downstream prioritization. The overall objective of this stage was to identify parasite proteins associated with essential biological processes, enriched metabolic functions, and potential therapeutic relevance, while also considering their divergence from the human host proteome.

Gene Ontology (GO) enrichment analysis was performed focusing on the Biological Process (BP) category using the *clusterProfiler* [22] and *enrichplot* [23] packages. Because organism-specific annotation resources for *T. cruzi* are limited compared with model organisms, an SQLite-based annotation package was generated for *T. cruzi* (NCBI Taxonomy ID: 353153) using the *AnnotationForge* package [24]. This custom annotation resource enabled GO term mapping, enrichment testing, and visualization of overrepresented biological processes within the analyzed proteome.

Enrichment results were filtered and organized to prioritize biological processes associated with parasite viability, metabolism, intracellular organization, and host–parasite adaptation. Particular attention was given to proteins involved in central metabolic pathways, energy production, macromolecule processing, and conserved biological processes that could represent vulnerabilities for pharmacological intervention. GO enrichment outputs were visualized using dot plots, bar plots, and enrichment network representations to identify functionally related clusters and to guide target prioritization.

To complement functional enrichment with network-based prioritization, protein–protein interaction (PPI) networks were retrieved from the STRING database (v12.0; https://string-db.org; accessed 30 April 2025) [25]. The resulting interaction networks were imported into Cytoscape [26] for topological and modular analysis. Molecular Complex Detection (MCODE) [27] was applied to identify highly interconnected regions that may represent functional modules or protein complexes. Together, network centrality analysis was performed using the cytoHubba plugin [28], ranking nodes according to the Maximal Clique Centrality (MCC) algorithm. Proteins with high MCC scores were considered potential hub proteins, as their centrality suggested functional importance within the parasite interaction network.

Functional annotation of the resulting network modules was further examined using the stringApp plugin in Cytoscape for Kyoto Encyclopedia of Genes and Genomes (KEGG) pathway enrichment [29]. This additional layer of analysis allowed the association of prioritized proteins with metabolic and regulatory pathways relevant to parasite survival. Proteins that were simultaneously represented in enriched GO categories, connected within PPI modules, and associated with essential metabolic pathways were considered stronger candidates for downstream structural and druggability evaluation.

To evaluate host–parasite selectivity, sequence similarity searches were performed against the human proteome (*Homo sapiens*, taxid:9606) using BLASTp [30]. For the identification of homologous sequences, an e-value threshold lower than 0.005 and a bit score higher than 100.0 were applied [31,32]. Proteins showing significant similarity to human proteins under these criteria were considered less favorable for selective drug discovery, whereas parasite proteins with limited or no significant human similarity were prioritized as potential selective targets. BLASTp similarity relationships between parasite proteins and human homologs were visualized using chord diagrams generated with the *circlize* package [33].

## Phylogenetic and Structural Analysis

To investigate the evolutionary conservation of the prioritized target, FASTA sequences corresponding to phosphopyruvate hydratase/enolase homologs were retrieved from the UniProt database. The *T. cruzi* selected sequences were used as the reference query, and sequence similarity searches were performed using BLASTp against the non-redundant protein database (https://www.nlm.nih.gov/ncbi/workshops/2023-08_BLAST_evol/databases.html, accessed 15 November 2024).

Multiple sequence alignment was performed in MEGA X using the ClustalW algorithm [34]. The resulting alignment was inspected to evaluate conservation patterns across taxa, with particular attention to residues and regions associated with catalytic activity and putative ligand-binding sites. Phylogenetic relationships were reconstructed using the neighbor-joining method, and the resulting tree was visualized and annotated using the Interactive Tree of Life (iTOL) platform [35].

## Structural Modeling and Protein Preparation of PPH from *T. cruzi* and *Leishmania donovani*

The amino acid sequence of phosphopyruvate hydratase (PPH) from *T. cruzi* was retrieved from the UniProt database (UniProt ID: Q4DMY6) [25]. To extend the analysis to kinetoplastid parasites beyond *T. cruzi*, sequence similarity searches were performed against *L. donovani* protein entries using BLASTp [34]. Based on BLASTp similarity against *L. donovani* protein entries and functional annotation, the putative enolase sequence UniProt ID A0A3Q8IJR5 [36] was selected as the closest *Leishmania* candidate for comparative structural analysis. This sequence showed 28.5% identity to the *T. cruzi* PPH query (Q4DMY6), with a bit score of 429 and an E-value of 2.5 × 10⁻⁴².

Because no experimentally resolved structures directly corresponding to the selected *T. cruzi* Q4DMY6 and *L. donovani* A0A3Q8IJR5 sequences were available, initial three-dimensional models were obtained from the AlphaFold Protein Structure Database [37]. The raw AlphaFold structures were inspected in Schrödinger Maestro to evaluate global fold integrity, domain organization, and the spatial distribution of low-confidence regions. Particular attention was given to the putative catalytic region and surrounding cavities, since these regions were intended for subsequent binding-site detection and ligand docking.

Both protein models were imported into Schrödinger Maestro and processed using the Protein Preparation Wizard [38]. Bond orders were assigned, hydrogen atoms were added, and formal charges were corrected where necessary. Protonation states were predicted at physiological pH using Epik/PROPKA at pH 7.0 ± 2.0 [39]. Hydrogen-bonding networks were optimized by reorienting hydroxyl groups, amide groups of asparagine and glutamine, and histidine side chains when required. Missing side chains were rebuilt, when necessary, whereas large loop remodeling was avoided to prevent artificial alteration of the predicted fold and putative binding regions.

Prepared structures were subjected to restrained energy minimization using the OPLS4 force field [40]. Minimization was performed to remove steric clashes and optimize local geometry while preserving the overall AlphaFold-predicted fold. These prepared models were then used as starting structures for MD-based refinement.

## Molecular Dynamics–Based Refinement and Representative Frame Selection

MD simulations were carried out with the Desmond engine (Schrödinger, LLC) [41] to refine the AlphaFold-derived models and obtain dynamically relaxed conformations for structure-based studies. Each protein was solvated in an orthorhombic box of TIP3P [42] water with a minimum 10 Å buffer from the protein surface to the box boundaries. Systems were neutralized and adjusted to 0.15 M NaCl. The OPLS4 force field was used throughout.

A multistage relaxation protocol was applied before production dynamics. First, energy minimization was performed using a hybrid steepest descent/conjugate gradient scheme until convergence of the maximum force (default Desmond criteria), to remove steric clashes and optimize local geometry.

This was followed by restrained equilibration under the NVT ensemble (constant number of particles, volume, and temperature) at 300 K for 1.0 ns, applying harmonic positional restraints to the protein heavy atoms (10 kcal·mol⁻¹·Å⁻²) to allow solvent relaxation around a quasi-fixed protein scaffold.

To enhance conformational sampling of the AlphaFold models, a simulated annealing protocol was then applied under NVT conditions, consisting of a temperature ramp from 300 K to 500 K over 200 ps, followed by a hold at 500 K for 200 ps, and subsequent cooling back to 300 K over 200 ps (total annealing time 0.6 ns). During annealing, backbone restraints were maintained (5 kcal·mol⁻¹·Å⁻²) to preserve the global fold while allowing sidechain and loop rearrangements.

After annealing, the system was equilibrated under the NPT ensemble (constant number of particles, pressure, and temperature) at 309.65 K and 1 atm in two consecutive stages:

i. 1.0 ns with reduced positional restraints on the backbone (2 kcal·mol⁻¹·Å⁻²), and
ii. 1.0 ns with minimal restraints (1 kcal·mol⁻¹·Å⁻²) to allow further relaxation of the protein environment. A final short unrestrained NPT equilibration (0.5 ns) was performed to ensure density convergence and pressure stabilization before production dynamics.

For model refinement production, MD simulations were carried out for 500 ns under NPT conditions (309.65 K, 1 atm) without positional restraints. These extended apo trajectories were used exclusively for structural relaxation, stereochemical monitoring, and representative frame selection before binding-site prediction and docking. Temperature was controlled using the Nosé–Hoover thermostat [43], and pressure was controlled using the Martyna–Tobias–Klein barostat [44]. Long-range electrostatic interactions were treated with the particle mesh Ewald (PME) method [45], and covalent bonds involving hydrogen atoms were constrained using the SHAKE algorithm [46], allowing a 2 fs integration time step.

Trajectory analyses included backbone root mean square deviation (RMSD) to assess global structural stability and root mean square fluctuation (RMSF) to identify flexible regions. Secondary structure elements (SSE) were monitored to confirm preservation of the overall fold during the simulation.

Representative frame selection was based on the combined evaluation of RMSD stabilization, RMSF behavior, preservation of secondary structure elements, catalytic-region integrity, and stereochemical quality metrics obtained from Ramachandran and protein geometry analyses. For *T. cruzi* PPH, the approximately 150 ns region was selected because it preserved an accessible catalytic-region topology while preceding the progressive stereochemical drift observed in later apo frames. The selected frame was subsequently subjected to geometry refinement and used for SiteMap pocket detection [47], docking, MM-GBSA calculations, and ligand-bound MD simulations.

The stereochemical metrics used to support frame selection are summarized in Supplementary Table S1.

For the *L. donovani* PPH-like model (UniProt ID: A0A3Q8IJR5), the same MD protocol was applied. A representative frame at approximately 70 ns (frame 141) was selected based on RMSD/RMSF profile, fold preservation, cavity accessibility, and stereochemical inspection.

The selected frames for both species were subsequently used for binding site prediction (SiteMap), receptor grid generation, molecular docking, MM-GBSA calculations, and ligand-bound MD simulations.

## Binding Site Identification, Catalytic Reference Mapping, and Grid Generation

Binding site identification was performed on the MD-refined representative structures of PPH using the SiteMap module in Schrödinger Maestro (v12.8.117) [47]. For each system, the selected frames (150 ns and 70 ns, respectively) were used as input structures. SiteMap calculations were carried out using standard parameters, allowing identification of the top-ranked potential binding pockets based on SiteScore, enclosure, exposure, hydrophobic/hydrophilic balance, and hydrogen-bonding capability.

Multiple candidate pockets were initially identified and ranked according to SiteScore. Although some sites presented higher numerical scores (≈1.04), the final pocket selection was not based solely on ranking. Instead, selection criteria incorporated structural and functional considerations, including spatial accessibility, cavity geometry, and consistency with catalytically relevant regions described for homologous enolase enzymes. A pocket corresponding to SiteScore ≈0.989 (designated as Pocket 3) was ultimately selected for sequences, as it exhibited appropriate size, solvent exposure, and proximity to residues associated with catalytic activity, including polar and charged residues capable of stabilizing ligand binding.

To map the catalytic reference region in *T. cruzi* PPH, the crystallographic *Trypanosoma brucei* enolase structure 2PTY was aligned to the *T. cruzi* model by backbone-based structural superposition [48]. Because 2PTY contains PEP coordinated with divalent metal ions, the crystallographic Zn²⁺ ions were replaced with Mg²⁺ ions for catalytic-site modeling, approximating the physiologically relevant metal-coordination state of enolase.

Following superposition, the relative positioning of PEP and the metal ions was analyzed to assess spatial correspondence with the SiteMap-predicted pocket. This combined structural mapping allowed refinement of the binding site definition beyond purely geometric criteria, integrating functional information derived from homologous enzymatic complexes. The resulting pocket was therefore considered a catalytically relevant binding region suitable for docking and molecular dynamics studies.

For the *L. donovani* PPH-like model, the 2PTY-derived catalytic reference was used only for structural comparison and was not used to define docking constraints. Receptor grid generation was performed using the SiteMap-selected pocket from the MD-refined 70 ns structure.

Once the binding site was defined for each system, receptor grids were generated using the Glide Receptor Grid Generation module. Grid boxes were centered on the centroid of the selected SiteMap pocket, with dimensions adjusted to fully enclose the binding cavity while minimizing inclusion of non-relevant solvent-exposed regions. No positional, distance, or pharmacophore constraints were applied during grid generation, allowing unbiased exploration of ligand binding modes during docking simulations.

## Ligand Preparation and Dataset Curation

Natural product (NP) compounds were selected from the Latin American Natural Product Database (LANaPDB) [20], which compiles structurally diverse molecules derived from Latin American biodiversity and has been widely used as a source of bioactive chemical space for drug discovery. The dataset includes compounds with reported or predicted biological activity, providing a relevant starting point for antiparasitic screening.

Chemical structures were retrieved in Simplified Molecular Input Line Entry System (SMILES)[49] format and imported into Schrödinger Maestro (v12.8.117) for preparation. Ligand preparation was performed using the LigPrep module [39], which generates energetically minimized three-dimensional conformations suitable for docking.

Protonation states and tautomeric forms were assigned using Epik at a physiological pH range of 7.0 ± 2.0, considering the local chemical environment of ionizable groups. This step ensured that ligands were represented in biologically relevant ionization states under simulation conditions.

To account for stereochemical and conformational diversity, up to 32 stereoisomers per compound were generated where applicable. All structures were then subjected to geometry optimization using the OPLS4 force field to obtain energetically favorable conformations before docking calculations.

Ligand structures were further inspected to ensure chemical integrity, including correct valence states, the absence of structural artifacts, and consistent ring conformations. Duplicate entries and improperly defined structures were excluded from the dataset.

Commercially available analogues of the prioritized virtual hit aptosimon were identified using ChemMine Tools with the PubChem fingerprint similarity option and a similarity threshold >0.9, followed by filtering for commercial availability in the MolPort catalog. Hinokinin was selected from this analogue search for focused docking, molecular dynamics simulations, and *in vitro* evaluation.

The final set of prepared ligands was used for molecular docking, MM-GBSA binding energy calculations, and molecular dynamics (MD) simulations to evaluate binding stability and interaction patterns with the PPH target.

## Chemical Space and Dataset Analysis

To characterize the chemical space of the natural product dataset and ensure structural diversity, an exploratory chemoinformatic analysis was performed. Molecular descriptors, including molecular weight (MW), lipophilicity (clogP), hydrogen bond donors (HBD), and acceptors (HBA), were calculated based on Lipinski’s Rule-of-Five criteria [50].

Principal component analysis (PCA) was applied to the dataset to evaluate the distribution of compounds in multidimensional chemical space [51]. PCA reduces dimensionality by projecting molecular descriptors into orthogonal principal components, enabling visualization of structural diversity. Three-dimensional PCA plots were generated to compare the distribution of compounds from LANaPDB [20] with those from other natural product libraries, confirming broad coverage of chemical space and the absence of clustering bias toward specific scaffolds.

*In silico* ADME and drug-likeness properties of aptosimon and hinokinin were estimated using SwissADME and complementary prediction outputs [52]. The evaluated descriptors included physicochemical properties, predicted solubility, gastrointestinal absorption, blood–brain barrier permeability, P-glycoprotein substrate status, cytochrome P450 inhibition profile, Lipinski drug-likeness, and synthetic accessibility (Supplementary Table S4).

## Molecular Docking

Structure-based molecular docking was performed using the Glide module implemented in Schrödinger Maestro (v12.8.117) [53,54]. Receptor grids were generated from the binding sites identified by SiteMap analysis, using the centroids of the selected pockets as reference points. Grid box dimensions were adjusted to fully enclose the binding cavity while minimizing inclusion of irrelevant solvent-exposed regions.

Docking simulations were carried out using a hierarchical protocol. Initially, all prepared ligands were screened using the Standard Precision (SP) mode to identify plausible binding poses and reduce the dataset size. Subsequently, top-ranked compounds from SP docking were subjected to Extra Precision (XP) docking for refined scoring and pose discrimination.

During docking, enhanced sampling was enabled, and a van der Waals scaling factor of 0.80 was applied to nonpolar ligand atoms to account for minor steric flexibility and improve the identification of energetically favorable binding conformations. Up to 10 poses per ligand were generated during SP docking, from which the top-ranked poses based on GlideScore were retained for XP refinement.

Docking results were ranked according to GlideScore, which approximates binding affinity by combining contributions from hydrophobic interactions, hydrogen bonding, electrostatics, and desolvation effects. Compounds were prioritized based on:

i. favorable GlideScore values relative to the dataset distribution,
ii. consistency of binding poses across SP and XP docking stages, and
iii. presence of interactions with residues located within the predicted binding pocket.

In addition to numerical scoring, visual inspection of docking poses was performed to evaluate the plausibility of ligand orientation, interaction patterns, and spatial complementarity with the binding cavity. Special attention was given to hydrogen bonding networks, π–π stacking interactions, and hydrophobic contacts with residues identified during SiteMap analysis.

Docking scores were normalized using Z-score transformation to evaluate the relative performance of each compound within the screened dataset. The resulting distributions were visualized using violin plots.

Selected ligands were further subjected to MM-GBSA binding free energy calculations and molecular dynamics simulations to evaluate the stability of the predicted complexes beyond static docking approximations.

For the selected compounds, the Tanimoto similarity score was calculated for clustering. The atom-pair-based fingerprints of the compounds were obtained using the “ChemmineR” package [55] in the R programming environment (version 4.0.3), and heatmaps were generated for visualization. Also, to examine the structural nature and similarity of the compounds that make up the dataset, a chemical diversity plot based on a substructure fragment dictionary-based binary fingerprint (FragFp) was also created using Osiris DataWarrior v05.02.01 software 61.

## MM-GBSA Binding Free Energy Calculations

Binding free energy estimations were performed using the Prime MM-GBSA module implemented in Schrödinger Maestro (v12.8.117) [56]. Calculations were carried out using the OPLS4 force field in combination with the VSGB 2.0 implicit solvation model [57], which accounts for solvent polarization effects and desolvation contributions upon ligand binding.

For each selected ligand, MM-GBSA calculations were initially performed on the minimized docking poses obtained from Glide XP. The binding free energy (ΔG_bind_) was computed according to the standard thermodynamic cycle:

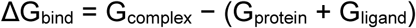

where G_complex_ corresponds to the free energy of the protein–ligand complex, and G_protein_ and G_ligand_ represent the energies of the isolated protein and ligand, respectively, calculated under identical conditions.

To incorporate dynamic information beyond static docking, additional MM-GBSA calculations were performed on representative frames extracted from equilibrated portions of the molecular dynamics trajectories. Frames were selected from regions exhibiting RMSD stabilization to ensure that energy estimations reflected thermodynamically relevant conformations.

For consistency across ligands, MM-GBSA calculations used receptor–ligand complexes derived from the same binding site definition identified through SiteMap. Ligands were evaluated under comparable conditions, and relative ΔG_bind_ values were used for ranking rather than absolute energy interpretation.

Particular care was taken when interpreting systems involving endogenous substrates and metal cofactors. Because phosphoenolpyruvate (PEP) is highly charged and physiologically coordinated by divalent metal ions, MM-GBSA values obtained without explicit metal coordination were considered non-comparable with those of neutral or weakly polar ligands. Therefore, PEP MM-GBSA results were interpreted qualitatively, whereas relative ΔG_bind_ comparisons were restricted to small-molecule ligands evaluated under consistent metal-free conditions, such as hinokinin and benznidazole.

MM-GBSA results were interpreted in conjunction with docking scores and molecular dynamics analyses, emphasizing consistency between energetic predictions and observed binding stability. Compounds showing favorable ΔG_bind_ values and stable interaction patterns during MD simulations were prioritized for further investigation and experimental validation.

## Molecular Dynamics Simulations of Ligand–Protein Complexes

MD simulations of ligand–protein complexes were performed using the Desmond MD engine (Schrödinger, LLC) [58] following the same general setup used for protein refinement, excluding simulated annealing steps, which were applied only during the initial refinement of the AlphaFold-derived protein structures.

These ligand-bound simulations were performed independently from the 500 ns apo-refinement trajectories and were used only to assess ligand stability, binding-mode persistence, and interaction dynamics.

Simulations were carried out for both *Trypanosoma cruzi* PPH and the *L. donovani* PPH-like model, using the representative frames selected from prior protein-only MD simulations (150 ns for *T. cruzi* and 70 ns for *L. donovani*). Protein–ligand complexes were constructed from Glide XP docking poses [53].

For *T. cruzi*, ligand systems included phosphoenolpyruvate (PEP) and benznidazole (BZ). For *L. donovani*, ligand systems included PEP and amphotericin B (AmB).

In the case of *T. cruzi*, additional systems incorporating divalent metal ions were prepared to evaluate catalytic-site configurations. Metal placement was guided by structural superposition with the crystallographic enolase from *Trypanosoma brucei* (PDB ID: 2PTY), where Zn²⁺ ions were replaced with Mg²⁺ ions to represent physiologically relevant cofactors. Ligand and metal positioning were defined before solvation based on the aligned structures.

All complexes were solvated in an explicit TIP3P water model within an orthorhombic simulation box, maintaining a minimum buffer distance of 10 Å between the protein and box boundaries. Systems were neutralized with counterions and adjusted to a physiological ionic strength of 0.15 M NaCl. The OPLS4 force field was applied to all systems.

Before production simulations, each system underwent a multistep relaxation protocol consisting of four steps, namely: (i) energy minimization, (ii) restrained NVT equilibration at 300 K for 1.0 ns with harmonic restraints applied to the protein backbone and ligand heavy atoms (10 kcal·mol⁻¹·Å⁻²), (iii) NPT equilibration at 309.65 K and 1 atm for 1.0 ns with reduced restraints (2 kcal·mol⁻¹·Å⁻²), and (iv) an additional 1.0 ns equilibration stage with minimal restraints (1 kcal·mol⁻¹·Å⁻²).

Production MD simulations were then performed under NPT conditions (309.65 K, 1 atm) for 300 ns without positional restraints. Temperature was maintained using the Nose–Hoover thermostat, and pressure was controlled using the Martyna–Tobias–Klein barostat. Long-range electrostatics were treated with the particle mesh Ewald (PME) method, and a 2 fs time step was used with bond constraints applied to hydrogen atoms.

Trajectories were recorded at regular intervals for analysis. Structural stability was evaluated using root mean square deviation (RMSD) of protein backbone and ligand heavy atoms, while residue-level flexibility was assessed through root mean square fluctuation (RMSF). Protein–ligand interactions were monitored over time using interaction timeline analysis tools implemented in Schrödinger Maestro.

Identical simulation conditions were applied across all systems to ensure comparability between *T. cruzi* and *L. donovani* complexes.

A summary of apo-refinement and ligand-bound MD systems, simulation lengths, and representative frames is provided in Supplementary Table S2.

## Antiparasitic Activity Assays

### Anti-*T. cruzi* intracellular assay

The anti-*Trypanosoma cruzi* activity was evaluated using the Tulahuen LacZ clone C4, which expresses β-galactosidase as a viability reporter [59]. Parasites were maintained in RPMI-1640 medium supplemented with L-glutamine, HEPES buffer, NaHCO₃, fetal bovine serum, and penicillin–streptomycin at 37 °C.

One day before infection, Vero cells were seeded in 96-well plates at 1.26 × 10⁴ cells/well in a final volume of 100 µL of culture medium. After 24 h, cells were infected with 5 × 10⁴ parasites/well diluted in 50 µL of culture medium. After an additional 24 h of infection, test compounds dissolved in DMSO were added and incubated for 120 h at 37 °C [60].

Antitrypanosomal activity was determined using chlorophenol red-β-D-galactopyranoside (CPRG; Roche Applied Science) as substrate for parasite β-galactosidase. CPRG was added to each well and allowed to react for 4.5 h at 37 °C. Absorbance was measured at 570 nm using a microplate reader [61]. Benznidazole was used as a positive control, and DMSO-treated infected cells were used as the negative control. IC₅₀ values were calculated from dose–response curves. In parallel, cytotoxicity was evaluated in non-infected Vero cells exposed to the same concentration range of hinokinin under comparable culture conditions. The CC₅₀ value was estimated as the concentration required to reduce Vero-cell viability by 50% relative to DMSO-treated controls and was used to calculate apparent selectivity indices as SI=CC₅₀/IC₅₀ [60].

### Anti-*Leishmania donovani* Promastigote Assay

The anti-*Leishmania* activity was evaluated against *L. donovani* promastigotes following the protocol described by Calderón et al. [62], using the fluorescent DNA intercalator PicoGreen (Invitrogen, USA). This assay was used as an initial parasite-growth inhibition screen for the selected compound.

For each bioassay, 1 × 10⁶ promastigotes/well were placed in 96-well plates with test compounds diluted in DMSO, in a final volume of 100 µL, and incubated for 3 days. Amphotericin B was used as the positive control [63].

After incubation, a PicoGreen cocktail was added at a 1:4 dilution and incubated at room temperature for 5 min. Fluorescence was then measured at 485 nm. Antileishmanial activity was expressed as percent inhibition of growth relative to the DMSO-treated negative control. IC₅₀ values were calculated from serial dilution assays.

For hinokinin, serial dilutions of 50, 25, 12.5, 6.25, and 3.125 µg/mL were tested in the antiparasitic assays and in the Vero-cell cytotoxicity assay. IC₅₀ and CC₅₀ values were estimated from dose–response data and expressed as the concentrations required to reduce parasite growth or Vero-cell viability by 50%, respectively. Apparent selectivity indices were calculated as SI = CC₅₀/IC₅₀, using the Vero-cell CC₅₀ and the corresponding antiparasitic IC₅₀ values.

## Results and Discussion

Figure 1 summarizes the complete methodological workflow, from proteome mining and target prioritization to structural refinement, ligand screening, MD simulations, and experimental evaluation of selected natural products.

### Proteome Mining, Functional Enrichment, and Target Prioritization

The proteome-wide analysis of *T. cruzi* identified a functionally diverse set of biological processes potentially relevant for parasite survival, adaptation, and therapeutic intervention. Functional enrichment analysis highlighted biological process categories associated with RNA modification, lipid metabolism, carbohydrate metabolism, intracellular organization, microtubule-associated processes, intracellular signaling, metabolic regulation, and overlapping functional categories associated with parasite adaptation and viability (Figure 2A-C). These enriched categories collectively indicate that *T. cruzi* depends on tightly coordinated metabolic and regulatory systems that support translational fidelity, membrane remodeling, intracellular adaptation, and energetic plasticity throughout its complex life cycle [64]. Rather than representing isolated pathways, the enrichment profile suggests the existence of highly interconnected biological networks required for parasite survival across distinct environmental and host-associated conditions. This system-level organization is notable from a therapeutic perspective because disruption of central metabolic hubs may propagate downstream effects across multiple adaptive pathways simultaneously.

**Figure 1.**
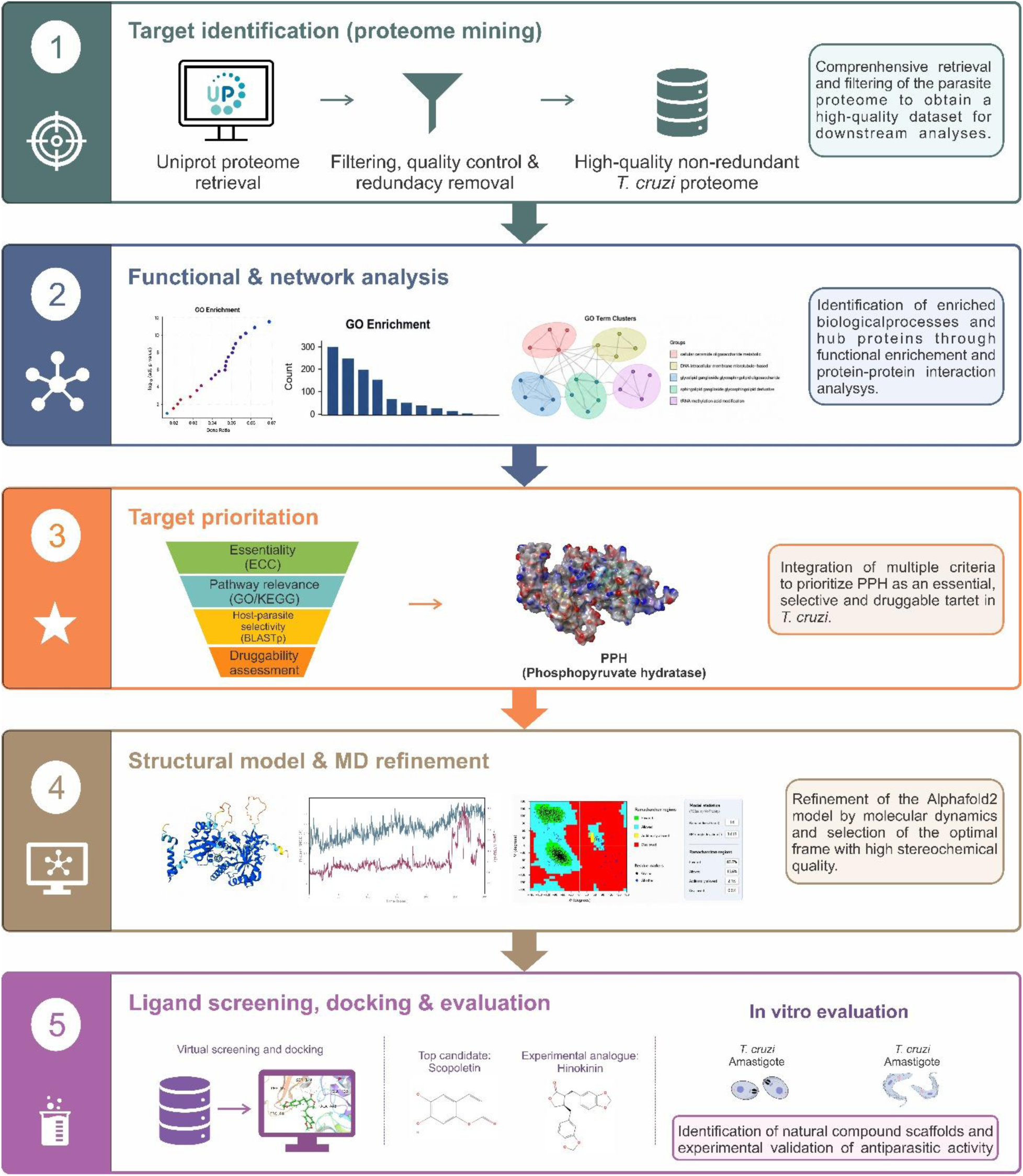
Integrated workflow for proteome-guided target prioritization, structural refinement, ligand screening, and experimental validation. The study integrated *T. cruzi* proteome mining, functional and network analysis, PPH prioritization, AlphaFold-based structural modeling, apo MD refinement, binding-site identification, virtual screening, ligand-bound MD simulations, and *in vitro* testing using *T. cruzi* and *Leishmania donovani* assays.

**Figure 2.**
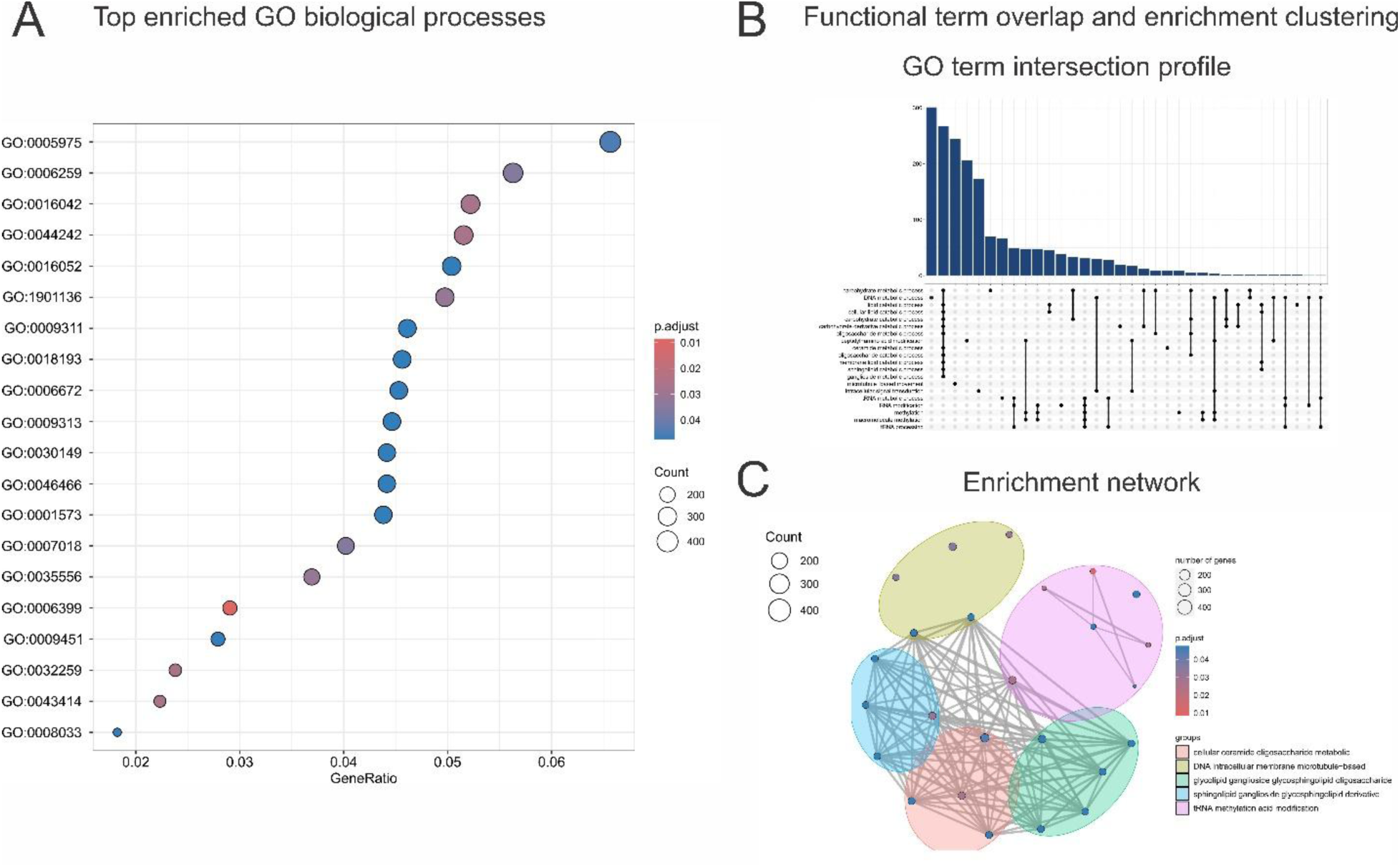
Functional enrichment analysis of *T. cruzi* proteins. (A) Dot plot highlights top Gene Ontology terms by gene ratio and adjusted p-value. (B) The bar and upset plots (top) show the distribution and overlap of enriched biological processes. (C) The network plot (bottom) groups related pathways emphasize key clusters in tRNA modification, lipid metabolism, and microtubule organization.

The bar and upset plots further demonstrated that several enriched biological processes were partially interconnected rather than functionally isolated, supporting the existence of coordinated metabolic and regulatory modules associated with parasite survival and adaptation (Figure 2B).

Among the enriched biological processes, RNA and tRNA modification-related categories were particularly relevant. Post-transcriptional regulation is central to kinetoplastid biology, as these organisms rely heavily on RNA processing and transcript stability rather than classical transcriptional regulation [65]. Thus, the enrichment of tRNA methylation and processing pathways suggests that maintenance of translational accuracy and proteostasis may represent a major vulnerability in *T. cruzi* [66]. Disruption of these processes could compromise parasite adaptation under host-derived stress conditions and impair stage-specific differentiation. Given the strong dependence of kinetoplastids on post-transcriptional regulation, these pathways may represent particularly attractive targets for selective therapeutic intervention [67,68].

Lipid-associated categories, including sphingolipid and glycosphingolipid metabolism, were also enriched. These pathways are biologically meaningful because lipid remodeling is required for membrane integrity, intracellular trafficking, parasite differentiation, and host–parasite interactions [69]. The enrichment of these pathways further supports the idea that membrane composition and lipid remodeling are not merely structural requirements but active components of parasite adaptation and persistence within the host environment. In addition, the enrichment of carbohydrate and oligosaccharide metabolism-related terms supports the importance of metabolic plasticity and energy homeostasis in parasite persistence [70]. Together, these results indicate that *T. cruzi* survival depends on a network of interconnected metabolic and regulatory processes rather than on a single isolated pathway [71].

Enrichment of microtubule-associated and intracellular signaling processes further indicated that structural organization, motility, cell division, and intracellular trafficking are also relevant components of *T. cruzi* adaptation (Figure 2C) [72]. These categories complement the metabolic enrichment profile by highlighting non-metabolic processes required for parasite survival, replication, and host interaction.

To refine target prioritization beyond enrichment analysis alone, the enriched proteome was further evaluated using protein–protein interaction network analysis and host similarity filtering (Figure 3). Network clustering identified groups of functionally connected proteins involved in parasite metabolism and cellular organization, while centrality analysis highlighted proteins with potential hub-like behavior within the interaction network [73]. The modular organization observed in the global PPI network suggests that prioritized proteins are distributed across coordinated functional groups rather than randomly connected nodes, supporting the existence of metabolically organized interaction modules associated with parasite viability (Figure 3A). This integrative network-based analysis was particularly important because proteins occupying central or highly connected positions may exert broader biological influence over parasite physiology [74]. In this context, highly connected metabolic proteins may represent strategic intervention points capable of simultaneously affecting multiple downstream pathways. However, because central metabolic enzymes are frequently conserved across eukaryotes, network relevance alone was insufficient for target prioritization [75]. Therefore, interaction topology was interpreted together with host similarity filtering and downstream structural tractability analyses.

**Figure 3.**
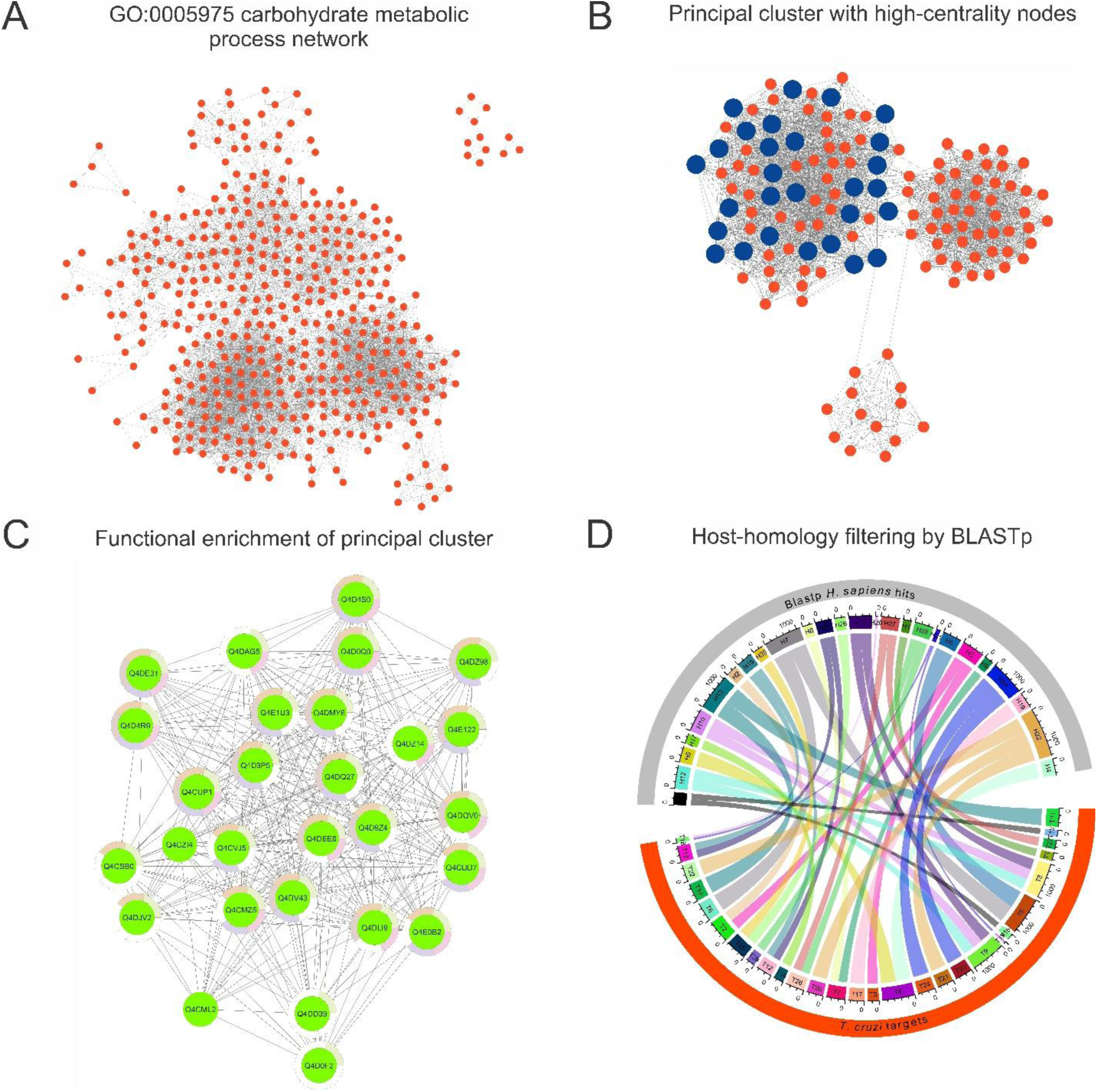
Refined network analysis of the carbohydrate metabolic process GO:0005975. (A) Protein–protein interaction network associated with GO:0005975, selected from the enrichment analysis shown in Figure 2. (B) Principal interaction cluster extracted from this network, with high-priority/high-centrality nodes highlighted in orange. (C) Functional enrichment analysis of the highlighted nodes, showing pathway categories linked to central metabolism, including glycolysis/gluconeogenesis, carbon metabolism, and the pentose phosphate pathway. (D) BLASTp-based comparison of prioritized parasite proteins against *Homo sapiens* homologs, supporting host-similarity filtering and prioritization of candidate targets, including phosphopyruvate hydratase (PPH) and phosphoglucomutase (PGM).

Functional enrichment of the highlighted nodes within this cluster showed strong association with metabolic pathways, biosynthesis of secondary metabolites, glycolysis/gluconeogenesis, carbon metabolism, and the pentose phosphate pathway (Figure 3C). These results supported further prioritization of enzymes within central carbon metabolism, including phosphopyruvate hydratase (PPH) and phosphoglucomutase (PGM), as candidate targets for downstream host-homology filtering and structural evaluation.

To evaluate potential host cross-reactivity, the proteins prioritized from this carbohydrate-metabolism-associated module were compared against the human proteome using BLASTp. This analysis allowed identification of parasite candidates with limited similarity to human homologs [76,77], supporting the prioritization of PPH and PGM as metabolically relevant candidates with potentially exploitable divergence from host proteins (Figure 3D).

Together, the GO:0005975 network refinement, cluster enrichment, and BLASTp-based host-homology filtering supported the prioritization of metabolically relevant parasite candidates with potential therapeutic selectivity.

Within this framework, phosphopyruvate hydratase (PPH) (UniProt ID: Q4DMY6), also known as enolase, emerged as a prioritized candidate. PPH catalyzes the reversible conversion of 2-phosphoglycerate into phosphoenolpyruvate, a key step in glycolysis [78], placing it within an essential energy-producing pathway. Beyond its canonical metabolic function, enolase has also been associated with non-glycolytic or “moonlighting” roles in several pathogens, including participation in host–parasite interactions and surface-associated functions [79]. This multifunctional behavior increases the biological relevance of PPH, as its inhibition could potentially affect parasite energy metabolism and additional parasite-specific adaptive processes linked to infection [80–82]. This observation provided the rationale for extending the downstream structural and ligand-binding analyses to a *L. donovani* PPH-like model as a comparative kinetoplastid system. This extension was not based on epidemiological overlap, but on the conservation of PPH/enolase-like enzymes across trypanosomatid parasites and the opportunity to evaluate whether ligand-accommodation features identified in *T. cruzi* could retain relevance in a related parasite model. The phylogenetic distribution observed across kinetoplastids therefore supported a comparative cross-species analysis of PPH-like structural features and ligand-binding behavior (Figure 4A) [83] [84]. The biological, structural, and selection criteria supporting PPH prioritization, together with the rationale for including the comparative *L. donovani* PPH-like model, are summarized in Table 1.

**Figure 4.**
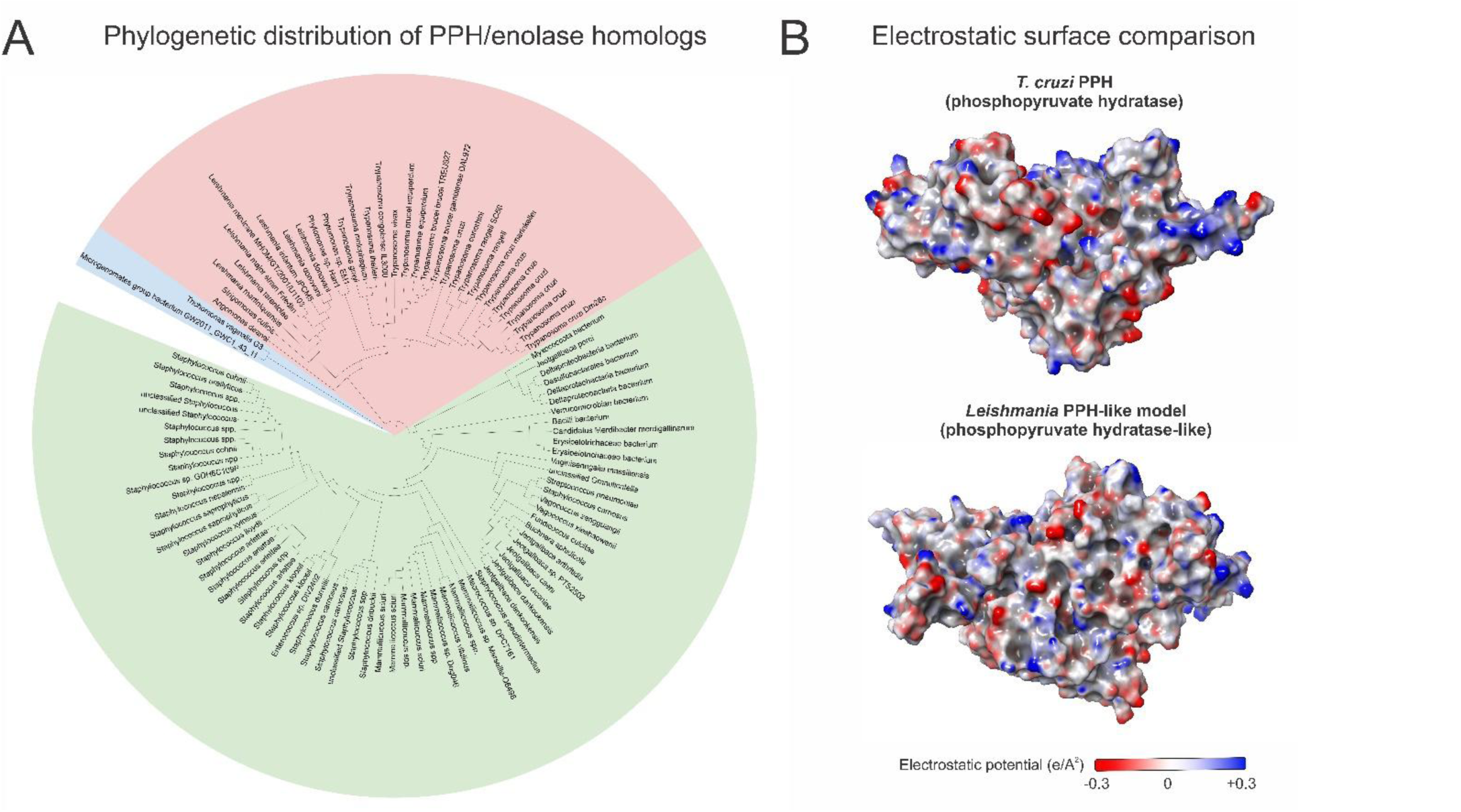
Phylogenetic distribution and electrostatic surface features of PPH-like enzymes. (A) Phylogenetic tree showing conservation of phosphopyruvate hydratase/enolase homologs across kinetoplastid parasites and broader eukaryotic taxa. Green sectors denote kinetoplastid species, whereas red sectors represent non-kinetoplastid eukaryotic groups. (B) Electrostatic surface potential maps of the MD-refined *T. cruzi* PPH model and the *Leishmania* PPH-like model, highlighting species-dependent charge distribution around surface-accessible regions used for downstream pocket and ligand-accommodation analyses.

**Table 1.**
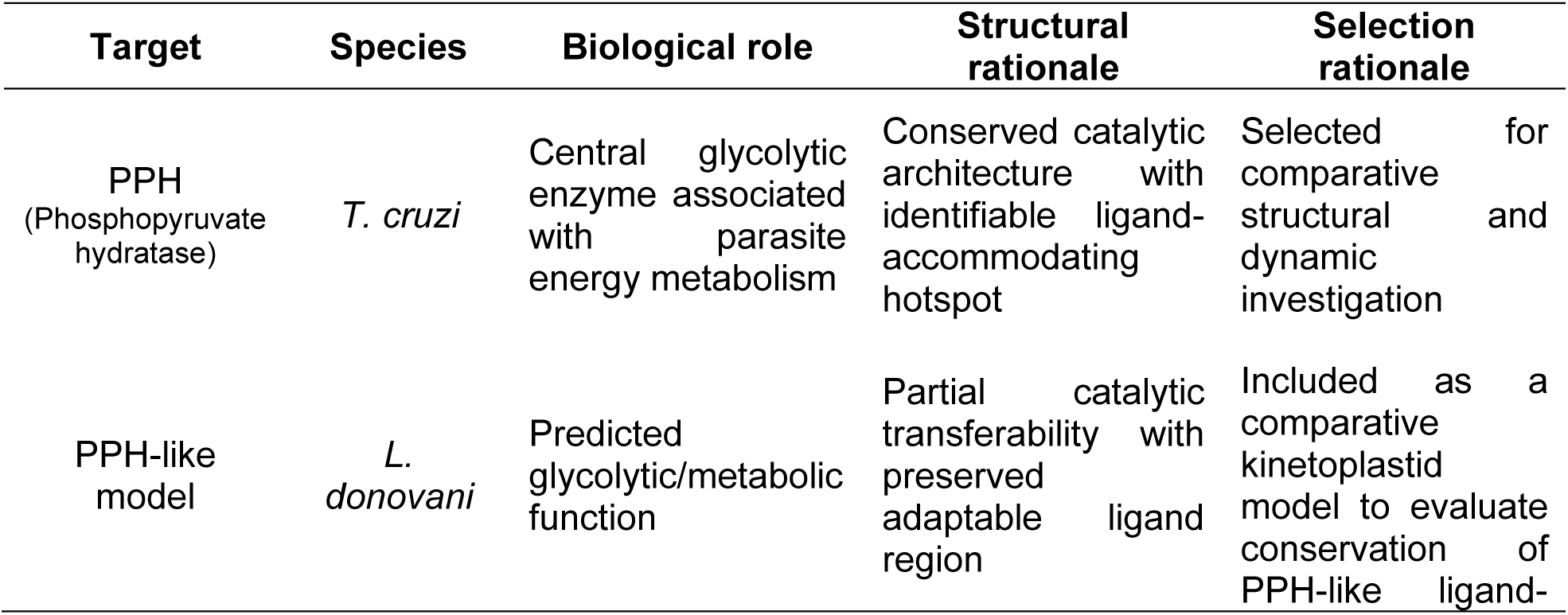
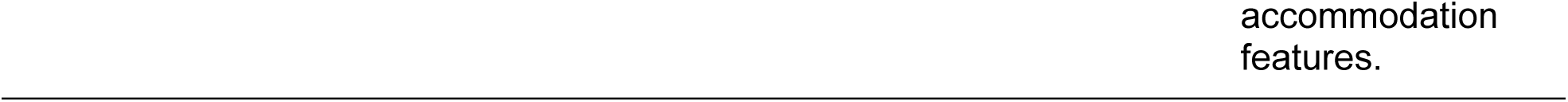
Rationale for PPH prioritization and comparative structural evaluation. Summary of the biological, structural, and selection criteria supporting the prioritization of *T. cruzi* PPH and the inclusion of a *L. donovani* PPH-like model for comparative ligand-accommodation analyses.

Structural conservation mapping further indicated that PPH contains a highly conserved catalytic core together with more variable peripheral regions (Figure 4B). This distinction is particularly important from a structure-based drug design perspective because conserved regions support catalytic functionality, whereas partially variable surface regions may provide opportunities for selective ligand recognition and species-adapted binding behavior [85]. Thus, Figure 4B highlights a structural rationale for subsequent pocket-centered analyses: highly conserved regions support catalytic relevance, whereas partially variable peripheral regions may provide opportunities for selective or species-adapted ligand accommodation.

At the same time, the broad conservation of PPH across kinetoplastids introduced an important structure-based challenge. Although conserved catalytic regions may favor the development of broad-spectrum antiparasitic inhibitors, they may also limit species selectivity if the catalytic architecture resembles host enzymes or remains highly preserved across related parasites. Hence, identifying structurally accessible and parasite-relevant ligand-accommodating regions—whether catalytic, adjacent to the catalytic core, or potentially allosteric—became essential for evaluating PPH as a selective therapeutic target [79].

Taken together, the enrichment, network, homology, and phylogenetic analyses converged on PPH as a biologically relevant and structurally tractable enzyme for downstream investigation. The detection of closely related PPH homologs in both *T. cruzi* and *L. donovani* further justified a comparative structure-based strategy aimed at determining whether natural-product-derived ligands could engage conserved or functionally analogous binding environments across kinetoplastid parasites. Importantly, the proteome-guided conservation patterns suggested that structurally relevant ligand-accommodating regions identified in *T. cruzi* might retain partial functional relevance across related kinetoplastids, supporting the rationale for extending the structural analyses beyond a single parasite species.

### Structural Model and Refinement

To ensure that downstream docking and ligand-bound MD simulations were performed on a structurally relaxed receptor, the AlphaFold-derived *T. cruzi* PPH model was subjected to extended apo MD refinement (Figure 1, step 4). Structural validation across trajectory frames showed that the approximately 150 ns region preserved the global fold and catalytic-region accessibility while avoiding the progressive stereochemical drift observed in later apo frames. The geometry-refined 150 ns structure was therefore selected as the representative receptor for SiteMap analysis, docking, MM-GBSA calculations, and ligand-bound MD simulations. Detailed Ramachandran comparisons and stereochemical metrics supporting this selection are provided in Supplementary Figure S1 and Supplementary Table S1.

### Chemical Space Characterization and Structure-Based Screening of Natural Products

Following proteome-guided target prioritization, a structure-based screening strategy was implemented to evaluate whether chemically diverse natural products from the LANaPDB could interact with structurally relevant regions of PPH (Figure 1, step 5). The workflow integrated chemical space analysis, ligand prioritization, binding-site characterization, docking evaluation, and molecular dynamics refinement to identify natural-product-derived scaffolds compatible with the enzyme’s predicted ligand-accommodating regions.

Following MD refinement, binding site analysis of the *T. cruzi* PPH model highlighted multiple surface-accessible cavities with characteristics compatible with small-molecule recognition. SiteMap analysis identified several candidate pockets with SiteScore values close to or above the commonly accepted druggability threshold (∼1.0). However, pocket selection was not based exclusively on numerical ranking. Instead, geometric accessibility, proximity to catalytically relevant residues, and consistency with the reference enolase structure from *T. brucei* (PDB ID: 2PTY) were collectively considered to define the most biologically plausible binding region.

Among the detected cavities, the selected pocket (Pocket 3; SiteScore ≈0.989) showed a favorable balance between enclosure, solvent accessibility, and spatial compatibility with the putative catalytic region (Figure 6A). Although other pockets exhibited slightly higher SiteScore values, these cavities were either excessively solvent-exposed or geometrically inconsistent with the catalytic architecture inferred from the comparative structural analysis. The selected pocket represented a compromise between predicted druggability and biological plausibility.

Structural superposition with the crystallographic *T. brucei* enolase complex revealed that the selected cavity partially overlapped with the substrate-associated region occupied by phosphoenolpyruvate (PEP) and divalent metal ions in the reference structure [48]. This overlap reinforced the functional relevance of the selected binding site and supported its use for docking and molecular dynamics analyses. Replacement of crystallographic Zn²⁺ ions with Mg²⁺ ions allowed construction of catalytic-like configurations compatible with the known metal dependence of enolase enzymes [83]. Visually, the resulting structural arrangement suggested the coexistence of a polar catalytic region associated with PEP/Mg²⁺ coordination and an adjacent mixed hydrophobic–polar cavity capable of accommodating non-substrate-like ligands (Figure 6A–C).

Principal component analysis (PCA) of the screened compounds revealed a broad and heterogeneous distribution of chemical space across the different LANaPDB-associated sources, including NuBBE_DB_, BIOFACQUIM, SistematX, UEFS, NAPRORE, UNIIQUIM, NPDB EjeCol, CIFPMA, PeruNPDB, and LAIPNUDELSAV (Figure 5A). The observed dispersion indicates that the dataset encompasses structurally diverse natural products rather than highly redundant chemical families, an important feature for exploratory drug discovery. Compounds derived from databases occupied partially overlapping but distinguishable regions of the PCA space, suggesting the coexistence of both shared and unique molecular scaffolds among regional biodiversity collections. This chemical heterogeneity is particularly advantageous in exploration and antiparasitic discovery because structurally diverse scaffolds may access distinct interaction regions within flexible enzymatic cavities.

**Figure 5.**
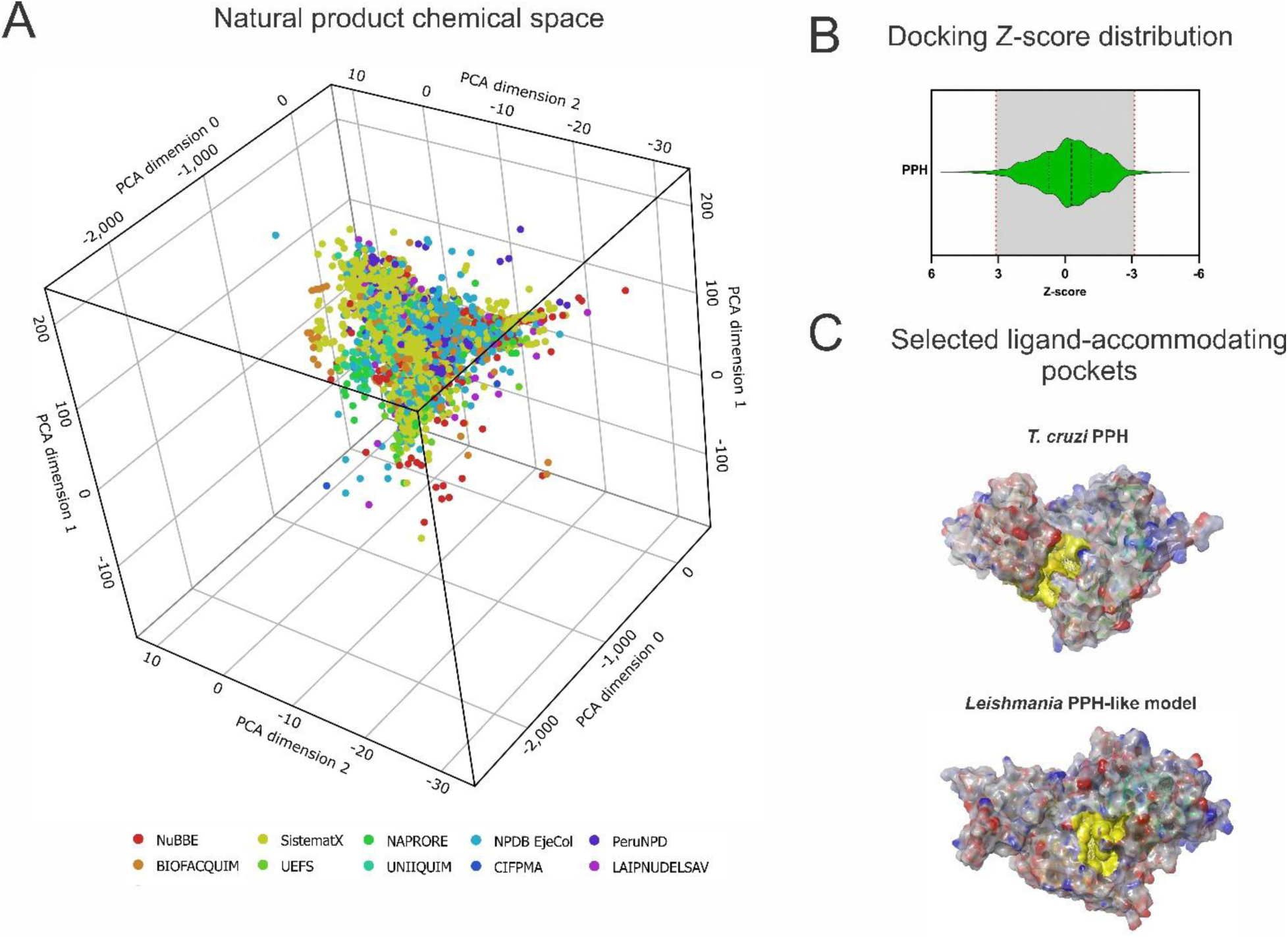
Chemical space distribution, docking-score prioritization, and pocket-centered screening strategy for PPH-like targets. (A) PCA-based chemical space distribution of the initial Latin American Natural Product Database included in the screening workflow. Colors indicate compound database source. (B) Docking Z-score distribution for compounds screened against PPH, highlighting the threshold-based prioritization region. (C) Electrostatic surface views of the MD-refined *T. cruzi* PPH and *L. donovani* PPH-like models, with SiteMap-selected ligand-accommodating pockets shown in yellow.

The PCA distribution also indicated that several compounds occupied regions compatible with drug-like physicochemical space, supporting their suitability for downstream docking analyses. Importantly, the absence of extreme clustering around a single chemical class suggested that the screening process was not dominated by a narrow scaffold family, reducing the likelihood of strong selection bias during virtual screening. This diversity is key in natural product-based drug discovery, where scaffold novelty and stereochemical complexity frequently contribute to unique interaction patterns with biological targets [86].

To evaluate the structural feasibility of ligand binding, the MD-refined PPH model was subjected to SiteMap analysis, revealing multiple cavities with characteristics compatible with small-molecule recognition (Figure 5C). Among the identified pockets, the selected region combined favorable geometric accessibility with proximity to catalytically relevant residues inferred from comparative structural analysis. The resulting binding environment presented a mixed polar and hydrophobic character, suggesting the potential accommodation of chemically diverse ligands [47].

Docking analyses and binding affinity estimations identified a subset of compounds with favorable interaction profiles against PPH. To facilitate comparison across the screened dataset, docking-derived scores were normalized using Z-score analysis after virtual screening (Figure 5B). A reference threshold of ±2.58 was used to highlight compounds located in the outer 1% of the docking-score distribution, whereas Z-score values beyond ±3 were interpreted as extreme outliers. Aptosimon (from *Homalomena wendlandii*) (LANaPDB10169) exceeded the more stringent Z-score threshold of 3 and defined an oxygenated lignan-like region of the prioritized set, supporting its selection as the original virtual hit for analogue-based follow-up [87]. Other high-ranking virtual hits, including Teuponin (from *Teucrium pernyi*) (LANaPDB4611) and Scopoletin (from *Amyris brenesii)* (LANaPDB692), also occupied favorable regions of the docking-score distribution. These compounds were retained as computationally prioritized scaffolds but were not advanced experimentally because the follow-up strategy required a commercially accessible analogue suitable for focused docking, molecular dynamics simulations, and *in vitro* validation. Aptosimon, therefore, served as the virtual hit, guiding the transition toward hinokinin as the experimentally evaluated lignan analogue.

Because aptosimon was not readily available for experimental testing, hinokinin was selected as a commercially available structural analogue using a PubChem fingerprint similarity threshold >0.9 and MolPort availability filtering as a source of commercially available compounds. Both compounds belong to an oxygenated lignan/lignan-lactone chemotype and retain benzodioxole-rich aromatic features that supported progression of hinokinin as a procurement-compatible representative of the prioritized chemical space [88,89]. *In silico* ADME and drug-likeness profiling further supported this progression, as both compounds showed oxygen-rich, drug-like physicochemical profiles, high predicted intestinal absorption, predicted blood–brain barrier permeability, and no predicted P-gp substrate liability. Relative to aptosimon, hinokinin showed slightly higher predicted lipophilicity and broader CYP inhibition liabilities, indicating that future optimization of this commercially accessible analogue should consider metabolic-interaction risk alongside antiparasitic potency (Supplementary Table S4).

To contextualize aptosimon within the broader prioritized docking subset, fingerprint-based similarity analysis was performed using FragFp descriptors (Supplementary Figure S2). This analysis was used to evaluate scaffold redundancy and chemical diversity, rather than to redefine the final experimental candidate. The similarity matrix and neighbor-network representation showed that the prioritized subset contained both closely related analogues and structurally divergent scaffolds. Similarity values approaching 1.0 identified closely related scaffold families, whereas lower values highlighted compounds that may support future analogue exploration and scaffold-hopping strategies.

To contextualize aptosimon within the broader prioritized docking subset, fingerprint-based similarity analysis was performed using FragFp descriptors (Supplementary Figure S2). This analysis was used to evaluate scaffold redundancy and chemical diversity, rather than to redefine the final experimental candidate. The similarity matrix and neighbor-network representation showed that the prioritized subset contained both closely related analogues and structurally divergent scaffolds. Similarity values approaching 1.0 identified closely related scaffold families, whereas lower values highlighted compounds that may support future analogue exploration, scaffold hopping, and alternative interaction patterns within the PPH binding cavity [90].

Overall, the combined chemical space, docking, similarity, and ADME analyses established a rational basis for progressing from the virtual hit aptosimon to the commercially available analogue natural product hinokinin for downstream MD simulations and biological validation.

### Binding site characterization and ligand interaction profiling in *T. cruzi* PPH

Docking analysis of PEP within the refined *T. cruzi* model produced a favorable GlideScore (−8.624 ± 0.171 kcal/mol), consistent with the expected affinity of a native substrate for the catalytic region. The docking pose positioned the phosphate-containing region of PEP near positively charged and polar residues lining the cavity [91], while also maintaining proximity to the Mg²⁺ coordination environment (Figure 6B). These observations support the structural consistency of the modeled catalytic region.

**Figure 6.**
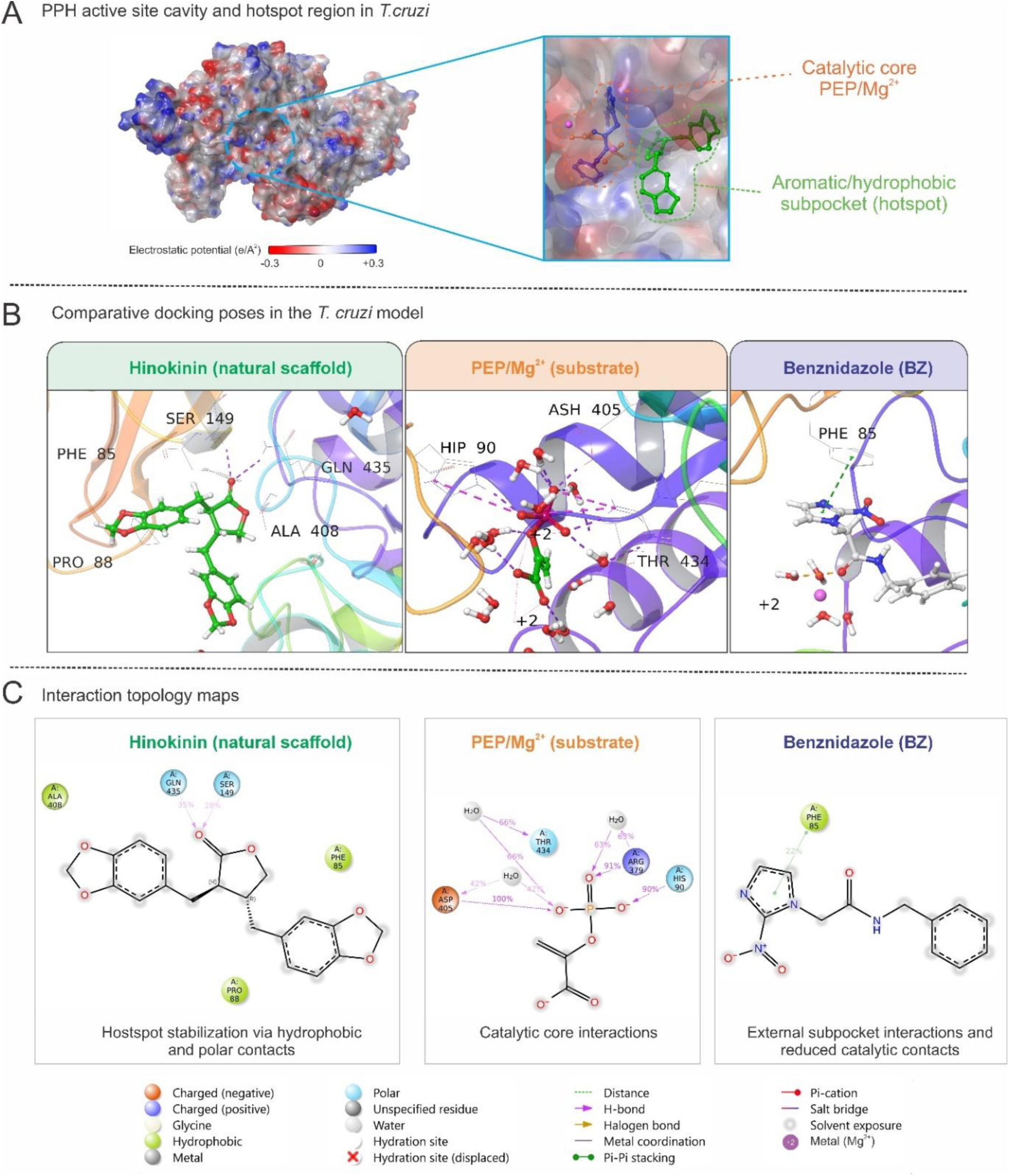
Binding-site characterization and ligand-accommodation behavior in *T. cruzi* PPH. (A) SiteMap-defined pocket and catalytic-reference mapping using the *T. brucei* enolase PEP/Mg²⁺ complex. (B) Comparative docking poses of PEP/Mg²⁺, hinokinin (aptosimon-like lignan analogue), and benznidazole within the selected cavity. (C) Two-dimensional interaction diagrams showing ligand-specific contact with the catalytic core and adjacent hotspot regions. Aptosimon was prioritized during virtual screening, whereas hinokinin was used as an experimentally accessible, structurally related scaffold for biological evaluation.

A comparative summary of hotspot persistence and catalytic-region behavior across ligand-bound systems is provided in Supplementary Table S3.

However, subsequent MM-GBSA calculations yielded highly unfavorable positive binding energy values for the PEP complex. This apparent discrepancy likely reflects the limitations of implicit-solvent MM-GBSA approaches when applied to highly charged substrate–metal systems. PEP is a strongly anionic metabolite whose interaction stability depends heavily on explicit electrostatic coordination with divalent metal ions and surrounding solvent molecules. Consequently, static MM-GBSA calculations performed under simplified conditions may not accurately reproduce the thermodynamic behavior of this type of catalytic complex. These observations also suggest that MM-GBSA estimations may provide more reliable comparative interpretation for neutral or weakly polar ligands than for highly charged substrate–metal assemblies whose stabilization depends strongly on explicit coordination dynamics.

Interestingly, although PEP yielded the most favorable GlideScore, the substrate-associated systems displayed unstable dynamic behavior during molecular dynamics simulations, including progressive displacement of Mg²⁺ ions and loss of the initial catalytic-like orientation. This behavior likely reflects the strong dependence of substrate stabilization on highly specific metal coordination and electrostatic organization, which may not be fully preserved in modeled systems lacking crystallographic catalytic constraints.

In contrast, the hinokinin-associated lignan analogue relied predominantly on mixed hydrophobic, aromatic, and water-mediated interactions rather than strict metal-dependent electrostatic coordination. This difference may explain why the natural-product-derived scaffold exhibited comparatively more persistent residence within the cavity despite more moderate docking scores.

To further evaluate the dynamic behavior of the catalytic configuration, MD simulations were performed for the PEP-bound system containing Mg²⁺ ions. During the trajectory, transient stabilization of substrate–metal coordination was observed during the early and intermediate stages of the simulation, followed by progressive displacement of the Mg²⁺ ions from the initially defined catalytic positions (Figure 7A). Despite this rearrangement, the overall protein fold remained stable throughout the simulation.

**Figure 7.**
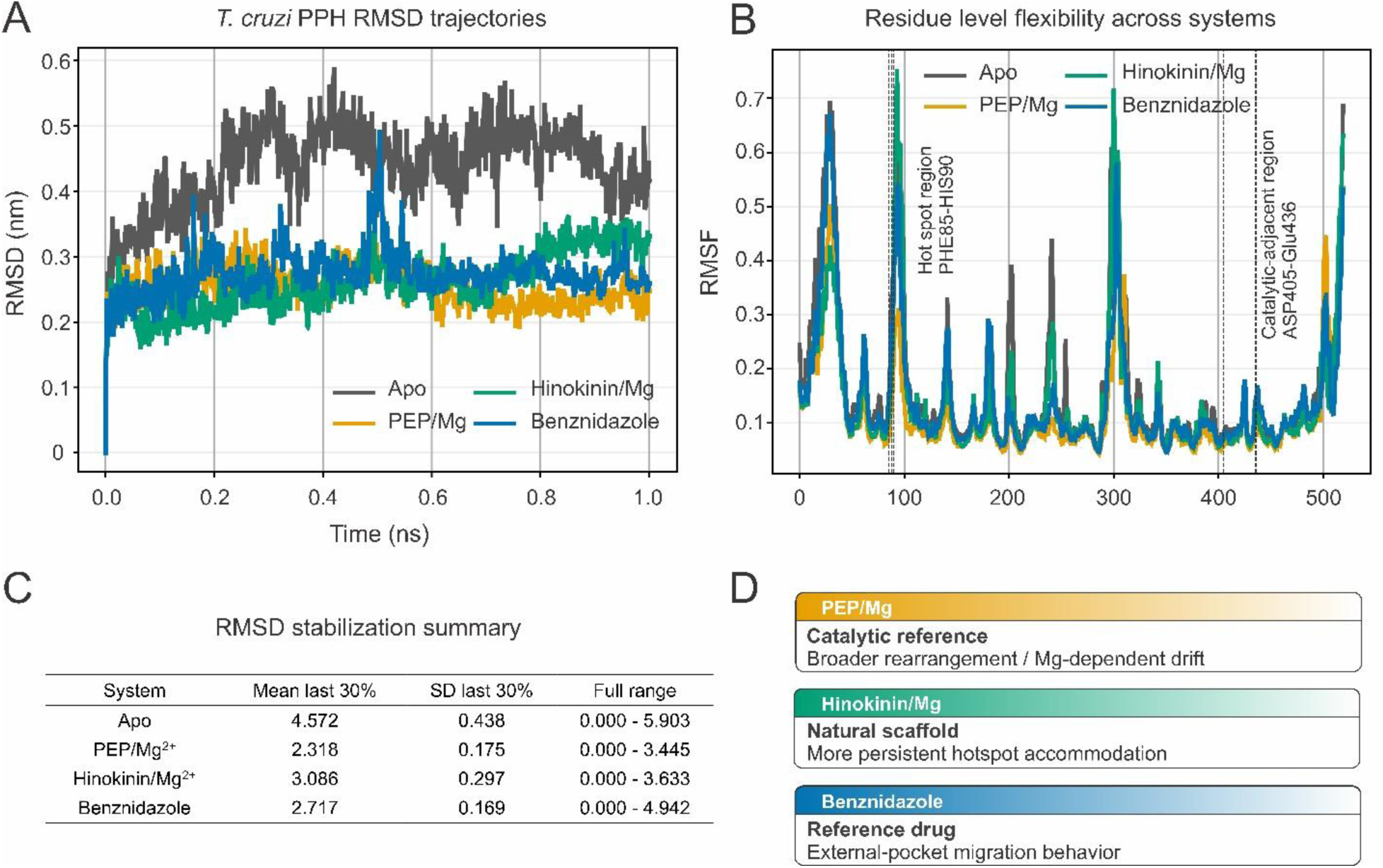
Ligand-bound molecular dynamic behavior of *T. cruzi* PPH complexes. (A) Backbone RMSD profiles of apo PPH and ligand-bound systems over 300 ns simulations, showing rapid stabilization of the hinokinin-associated complex and broader fluctuations in the PEP/Mg²⁺ system. (B) Ligand RMSD profiles showing ligand-dependent residence behavior within the selected cavity. (C) Residue-level RMSF profiles highlighting local flexibility changes across pocket-lining and catalytic-adjacent regions. (D) Comparative interpretation of ligand-dependent stabilization, showing persistent accommodation of the aptosimon-like lignan analogue and progressive rearrangement of the substrate-like PEP/Mg²⁺ configuration.

The observed mobility of Mg²⁺ ions may reflect the dynamic nature of metal coordination within the modeled catalytic environment rather than the simple instability of the protein fold itself. Importantly, the initial maintenance of substrate-associated interactions supports the functional plausibility of the selected cavity, even if long-timescale stabilization of the exact crystallographic configuration was not preserved under the simulated conditions.

To investigate the interaction behavior of screened ligands independently of explicit catalytic metal coordination, additional docking and MM-GBSA calculations were performed using the refined protein model without Mg²⁺ ions. Under these conditions, the hinokinin displayed a GlideScore of approximately −3.41 ± 0.181 kcal/mol and an MM-GBSA binding energy of approximately −24.15 ± 0.081 kcal/mol. Benznidazole exhibited a slightly more favorable docking score (approximately −4.77± 0.189 kcal/mol) and MM-GBSA value (approximately −28.05 ± 1.071 kcal/mol).

Although the absolute docking scores were moderate relative to the endogenous substrate, both ligands showed energetically favorable interaction profiles within the selected cavity. Importantly, ligand stability during molecular dynamics simulations did not strictly correlate with docking score magnitude alone, highlighting the importance of evaluating interaction persistence under dynamic conditions rather than relying exclusively on static docking estimations.

Analysis of ligand interaction patterns revealed that the selected *T. cruzi* PPH cavity contains both a catalytic polar subregion and an adjacent hydrophobic/aromatic subregion. In the PEP–Mg²⁺ system, ligand stabilization was dominated by persistent contacts with ASP405, ARG379, and HIS90, together with water-mediated interactions involving THR434, GLU436, and GLN435. These residues define a polar catalytic interaction network consistent with substrate-like recognition.

In contrast, hinokinin engaged a partially overlapping but more peripheral region, with recurrent interactions involving GLN435, SER149, PHE85, ALA408, and PRO88. Notably, PHE85 contributed hydrophobic and aromatic stabilization, whereas GLN435 provided recurrent polar contacts, suggesting that these residues may form part of a ligand-accommodating hotspot adjacent to the catalytic region. Benznidazole also sampled this region, sharing contacts with PHE85, PRO88, GLN435, and GLU436, although its interaction pattern was more transient and shifted toward a more external pocket during the trajectory.

The overlap of PHE85, GLN435, and GLU436 across hinokinin and benznidazole, together with the partial contribution of ASP405 and ARG379 in substrate-associated configurations, suggests that the selected cavity should not be interpreted as a single rigid catalytic pocket. Instead, it appears to behave as a composite binding region composed of a polar PEP/Mg²⁺-associated core and an adjacent hydrophobic/aromatic subpocket capable of accommodating non-substrate-like ligands.

Interaction persistence analysis suggested that residues PHE85, GLN435, and GLU436 behaved as recurrent ligand-accommodating nodes across multiple simulations, whereas catalytic residues such as ASP405 and ARG379 contributed predominantly to substrate-associated stabilization. This distinction supports the existence of a dynamically heterogeneous cavity containing partially specialized interaction subregions rather than a single rigid catalytic pocket.

Comparison with the crystallographic *T. brucei* enolase–PEP complex (PDB ID: 2PTY) provided a structural reference for interpreting the catalytic region of the *T. cruzi* PPH model. In 2PTY, PEP is surrounded by a highly polar and metal-coordinating environment involving residues such as SER373, ARG372, HIS371, LYS394, HIS156, GLN164, GLU165, ASP243, GLU291, and ASP318, with the crystallographic Zn²⁺ ions coordinated mainly by acidic residues. After structural superposition and replacement of Zn²⁺ by Mg²⁺, the modeled *T. cruzi* PEP–Mg²⁺ system reproduced the general requirement for a polar, metal-dependent substrate-recognition environment, although the exact residue numbering and local geometry differed between species.

MD simulations further identified distinct binding profiles between the evaluated ligands. Benznidazole initially maintained interactions near the catalytic region but progressively migrated toward a more external region of the cavity during later stages of the trajectory. In contrast, the hinokinin-associated lignan analogue displayed comparatively stable residence within the selected binding region over the simulation timeframe. These differences reinforce the notion that docking scores alone may not fully capture the dynamic stability of ligand binding within the PPH cavity.

RMSD analysis indicated that the hinokinin-associated system stabilized rapidly within the first 10–15 ns and fluctuated predominantly within a ∼0.35–0.60 nm range throughout the simulation, whereas the apo structure stabilized at higher average deviations (∼0.55–0.65 nm). In contrast, the PEP/Mg²⁺ system exhibited broader fluctuations and progressive destabilization of the catalytic arrangement during later simulation stages.

Residue-wise RMSF analysis revealed damped fluctuations around pocket-lining regions associated with ligand accommodation, including residues surrounding the aromatic stabilization region previously associated with Trp398-like interactions. These reduced fluctuations support persistent local stabilization of the binding environment throughout the trajectory.

Overall, the observed interaction patterns support the interpretation of the selected PPH region as a dynamically adaptive cavity capable of accommodating chemically distinct ligands through partially overlapping interaction networks rather than through a single rigid substrate-recognition geometry.

Collectively, these analyses support the structural relevance of the selected *T. cruzi* PPH binding region and demonstrate that the refined model can accommodate both endogenous substrate-like configurations and structurally distinct natural-product-derived scaffolds. The combined docking, MM-GBSA, and molecular dynamics results further emphasize the importance of integrating static and dynamic approaches when evaluating ligand behavior in modeled enzymatic systems.

### Comparative Structural and Ligand-binding Analyses in the *L. donovani* PPH-like Model

The conservation of phosphopyruvate hydratase across kinetoplastid parasites motivated extension of the structural and ligand-binding analyses to an *L. donovani* PPH-like model. The selected *L. donovani* sequence, A0A3Q8IJR5, was identified as the closest annotated enolase-like candidate to the *T. cruzi* PPH query by BLASTp, showing 28.5% identity, a bit score of 429, and an E-value of 2.5 × 10⁻⁴². Despite this moderate sequence identity, phylogenetic clustering and functional annotation supported its classification as a PPH/enolase-like homolog, while comparative molecular dynamics refinement revealed local differences in conformational flexibility and cavity organization that influenced subsequent docking and ligand-binding behavior.

The *L. donovani* model retained the characteristic α/β architecture observed in the *T. cruzi* structure, but exhibited increased mobility in loop-rich and surface-accessible regions during molecular dynamics simulations (Figure 8A). RMSD analysis demonstrated earlier local stabilization followed by gradual conformational drift at later stages of the trajectory, suggesting that the binding cavities of the *L. donovani* model remained more dynamically plastic throughout the simulation.

**Figure 8.**
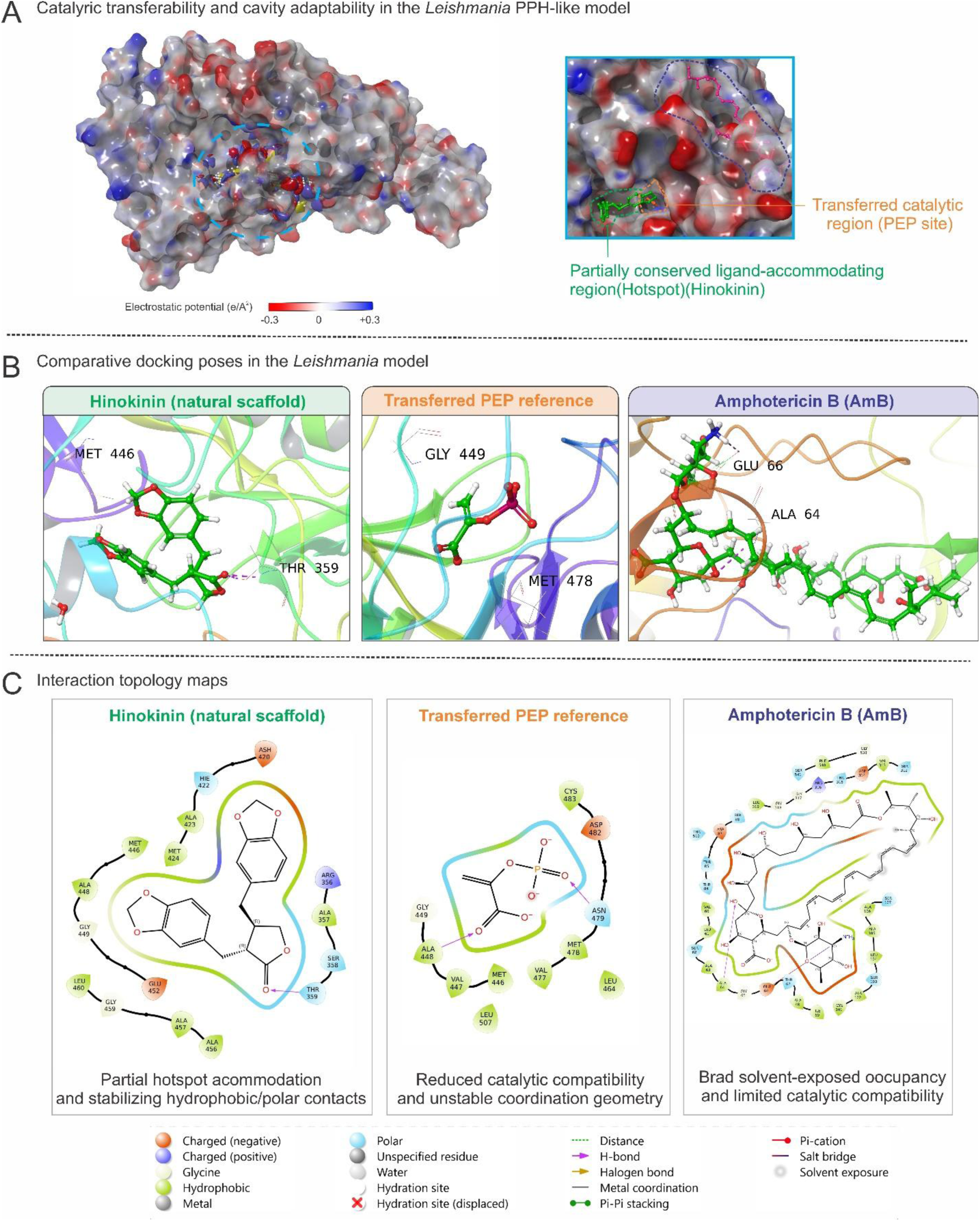
Comparative structural and ligand-accommodation behavior of the Leishmania PPH-like model. (A) MD-derived structural behavior of the Leishmania PPH-like model, highlighting increased flexibility in loop-rich and surface-accessible regions. (B) Structural mapping of the PEP/Mg²⁺ catalytic reference from T. brucei enolase onto the Leishmania model, showing limited transferability and partial burial of the substrate-like configuration. (C) Docking and interaction analyses showing favorable accommodation of hinokinin within a SiteMap-defined accessible cavity.

Despite these structural variations, hinokinin maintained favorable energetic behavior within the *L. donovani* PPH-like cavity, supporting the ability of this aptosimon-related lignan analogue to engage partially conserved ligand-accommodating regions across the evaluated kinetoplastid PPH models.

For this reason, a representative frame extracted at approximately 70 ns (frame 141) was selected for downstream analyses, as this region of the trajectory preserved accessible cavities before the larger structural rearrangements observed at later stages. The selected structure displayed a solvent-accessible binding environment compatible with small-molecule docking while maintaining the overall integrity of the enolase-like fold.

In contrast to the *T. cruzi* model, direct transfer of the catalytic configuration derived from the crystallographic *T. brucei* enolase structure (PDB ID: 2PTY) proved less structurally compatible with the *L. donovani* PPH-like model. Structural superposition revealed that the PEP molecule from the reference structure became partially buried within the modeled protein interior rather than occupying a clearly accessible catalytic cavity (Figure 8B). This observation suggested that local structural organization around the catalytic region differed sufficiently to limit direct extrapolation of the crystallographic substrate configuration.

Consequently, binding site identification in the *Leishmania* model relied primarily on SiteMap-predicted cavities derived directly from the MD-refined structure rather than on strict catalytic-site transfer from the reference enolase complex. The selected cavities exhibited mixed polar and hydrophobic characteristics similar to those observed in *T. cruzi*, although with distinct geometric organization and altered accessibility of surrounding regions.

Docking analyses demonstrated that the hinokinin, selected as an experimentally accessible natural-product scaffold related to the prioritized chemical space, maintained favorable interaction behavior within the *Leishmania* cavity. The ligand produced a GlideScore of approximately −5.588 ± 0.382 kcal/mol and an MM-GBSA binding energy of approximately −42.96 ± 0.875 kcal/mol, representing the most favorable energetic profile among the evaluated small molecules in the *Leishmania* system (Figure 8C). These results suggest that the identified cavity is compatible with stabilization of aromatic oxygenated scaffolds despite local structural differences relative to the *T. cruzi* model.

In contrast, docking of phosphoenolpyruvate produced a comparatively weaker GlideScore (approximately −3.125 ± 0.531 kcal/mol) and unfavorable MM-GBSA behavior. Similar to the *T. cruzi* catalytic model, these results likely reflect the limitations of simplified energetic calculations for highly charged substrate-like molecules whose stability depends strongly on explicit electrostatic coordination and solvent effects.

Amphotericin B (AmB), included as a reference antileishmanial compound, produced a favorable docking score (approximately −5.04 ± 0.357 kcal/mol) and strongly negative MM-GBSA binding energy (approximately −49.44 ± 0.114 kcal/mol). However, visual inspection of the docking pose revealed that the large macrolide structure occupied an extended region partially outside the predicted cavity (Figure 8B). The size and structural complexity of AmB suggested that its interaction behavior was not directly comparable with the smaller substrate-like ligands evaluated in the same cavity.

Because of this marked difference in molecular size and cavity occupancy, long-timescale molecular dynamics simulations were not pursued for the AmB-bound system. Instead, subsequent MD analyses focused primarily on the endogenous substrate-like configuration (PEP) and hinokinin, which displayed interaction geometries more compatible with the dimensions of the selected cavity.

MD simulations of the PEP-associated *L. donovani* system further reinforced the structural differences between the two kinetoplastid models. Unlike the partial catalytic stabilization observed transiently in *T. cruzi*, the substrate configuration in *L. donovani* rapidly lost the initial binding orientation during the early stages of the simulation, with displacement of the ligand away from the originally defined catalytic arrangement. These observations support the notion that the catalytic environment inferred from the *T. brucei* reference structure is less structurally transferable to the *Leishmania* PPH-like model.

Despite these differences, the persistence of favorable interaction behavior for hinokinin suggests that the selected cavity retains the capacity to accommodate non-substrate-like ligands across kinetoplastid species. This observation is significant because it indicates that ligand recognition may not require strict conservation of the canonical substrate-binding arrangement.

Notably, the interaction behavior of hinokinin did not fully reproduce the canonical substrate-binding geometry observed in crystallographic enolase systems. Instead, the ligand preferentially stabilized adjacent polar–hydrophobic regions surrounding the catalytic environment, suggesting the possibility of non-canonical ligand accommodation within the broader PPH cavity architecture.

Taken together, the comparative analyses demonstrate that although PPH-like proteins from *T. cruzi* and *L. donovani* share an overall conserved fold, local differences in cavity accessibility, flexibility, and catalytic-site organization influence ligand-binding behavior. These findings reinforce the importance of species-specific structural refinement and dynamic evaluation when extending structure-based drug discovery strategies across related parasitic systems.

Comparative analysis between the two kinetoplastid systems indicated that although overall fold conservation was preserved, substrate-associated catalytic configurations were substantially more transferable to the *T. cruzi* model than to the *L. donovani* structure. In contrast, hinokinin retained comparatively stable accommodation behavior in both systems, suggesting that non-substrate-like ligand recognition may be less sensitive to species-dependent catalytic rearrangements.

### Experimental Antiparasitic activity and correlation with structure-based analyses

To assess whether the chemical space prioritized from the virtual hit aptosimon retained biological activity in cellular systems, hinokinin was evaluated as the commercially accessible structural analogue selected for experimental testing. Hinokinin was tested against intracellular *T. cruzi* amastigotes and *L. donovani* promastigotes. The compound showed reproducible activity against *T. cruzi*, with IC₅₀ values of 3.90 and 3.52 ± 0.023 µg/mL in two independent determinations, corresponding to a mean IC₅₀ of approximately 3.7 µg/mL. Hinokinin also showed growth-inhibitory activity against *L. donovani* promastigotes, with an IC₅₀ of 13.06 ± 0.018 µg/mL. Using the Vero-cell CC₅₀ value of 36.89 ± 0.751 µg/mL, the apparent selectivity indices were approximately 9.9 for *T. cruzi* and 2.8 for *L. donovani* (Table 2).

**Table 2.**
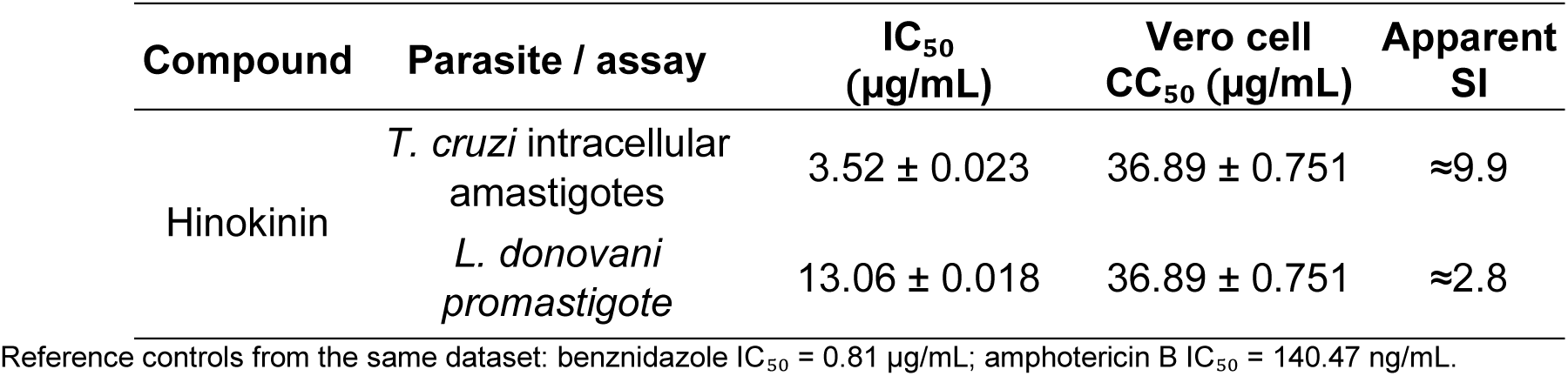
Antiparasitic activity and apparent selectivity of hinokinin. IC₅₀ values are shown for hinokinin against intracellular *T. cruzi* amastigotes and *L. donovani* promastigotes, together with the CC₅₀ value determined in non-infected Vero cells. Apparent selectivity indices were calculated as SI = CC₅₀/IC₅₀ using the Vero-cell CC₅₀ and the corresponding antiparasitic IC₅₀ values.

The activity observed against *T. cruzi* is consistent with previous reports supporting the trypanocidal potential of hinokinin, including a hinokinin-loaded microparticle formulation that significantly reduced parasitemia in infected mice [92]. Because that study evaluated a formulation rather than the free compound, the comparison should be considered supportive rather than directly equivalent.

Because the *L. donovani* assay was performed using promastigotes, this result should be interpreted as an initial antileishmanial growth-inhibition signal rather than as direct evidence of activity against the clinically relevant intracellular amastigote stage.

This activity range is relevant because hinokinin was not optimized through medicinal chemistry but was selected as the experimentally accessible analogue of the virtual hit aptosimon. Previous work has already described the antitrypanosomal potential of hinokinin and related lignan lactones [93], including activity against *T. cruzi* forms. Thus, the present results do not position hinokinin as a newly discovered antitrypanosomal molecule, but rather as an experimental validation layer for the target-guided structural workflow, linking the aptosimon-prioritized chemical space with hinokinin activity and PPH-centered ligand accommodation.

The biological activity observed experimentally was broadly consistent with the structure-based analyses performed using the *T. cruzi* PPH model. Although the absolute docking score for hinokinin was moderate, molecular dynamics simulations suggested comparatively persistent ligand accommodation within the selected cavity. This discrepancy between docking score magnitude and dynamic stability was also evident when comparing the evaluated ligands: phosphoenolpyruvate displayed highly favorable docking behavior within the catalytic region but failed to maintain stable substrate-associated configurations throughout long-timescale simulations, whereas hinokinin showed less extreme docking scores but more persistent interaction behavior during MD simulations. These observations support the use of docking, MM-GBSA, and molecular dynamics as complementary metrics rather than relying on a single static score.

Notably, the experimentally observed antiparasitic activity correlated more consistently with persistent ligand accommodation during molecular dynamics simulations than with absolute docking score magnitude alone. This observation reinforces the importance of incorporating dynamic structural evaluation into antiparasitic screening workflows involving flexible enzymatic systems.

Benznidazole, used as the reference antitrypanosomal drug [94], also displayed energetically favorable interaction behavior within the selected cavity. However, MD simulations suggested progressive migration of the molecule toward more external regions of the binding environment during later stages of the trajectory. Although the present study was not intended to reproduce the complete pharmacological mechanism of benznidazole, these observations suggest that the selected cavity may accommodate multiple ligand interaction modes rather than functioning as a rigid substrate-exclusive catalytic pocket.

The antileishmanial assay further supported the cross-species relevance of the selected chemotype. Hinokinin showed measurable activity against *L. donovani*, and the protein model retained favorable interaction behavior for hinokinin despite structural differences relative to the *T. cruzi* cavity. Together, these data suggest that non-substrate-like ligand accommodation may be conserved more readily than the canonical substrate configuration across kinetoplastid PPH homologs.

This observation is critical because the *Leishmania* model displayed reduced compatibility with direct transfer of the catalytic PPH configuration derived from the *T. brucei* reference structure. Nevertheless, the selected natural-product-derived scaffold remained compatible with the SiteMap-defined cavity, suggesting that ligand recognition may occur independently of strict substrate mimicry. Such behavior may represent an advantage for future lead optimization, as compounds would not necessarily need to reproduce the highly charged catalytic interactions associated with endogenous metabolites.

The favorable MM-GBSA result calculated for hinokinin further supports this interpretation. In contrast to PPH, whose energetic behavior remained strongly influenced by metal coordination and electrostatic complexity, the natural-product-derived scaffold exhibited more stable and interpretable energetic profiles under the evaluated conditions.

Amphotericin B, used as the positive control in antileishmanial assays [95], exhibited expected potent activity. However, computational analyses indicated that its large macrolide architecture occupied an extended region beyond the dimensions of the selected cavity. This suggests that the experimentally observed antileishmanial activity of amphotericin B is unlikely to depend on the same interaction environment explored for the smaller ligands in the present study. Consequently, the amphotericin B docking results were interpreted primarily as comparative structural references rather than direct mechanistic equivalents.

An important aspect emerging from the combined computational and experimental analyses is the apparent tolerance of the selected PPH cavities to chemically distinct scaffolds. The evaluated ligands included substrate-like molecules, nitroheterocyclic compounds, and large polyene macrolides, yet several retained measurable interactions and compatibilities with the modeled systems. This structural permissiveness may reflect the dynamic nature of the selected cavities and suggests the existence of adaptable interaction environments rather than highly rigid binding geometries.

The integrated workflow employed in this study—from proteome mining and phylogenetic conservation analysis to molecular dynamics refinement, docking, and experimental validation—therefore supports the biological relevance of PPH-like proteins as structure-based targets in kinetoplastid parasites. At the same time, the results emphasize that catalytic-site behavior in modeled enzymes cannot always be inferred directly from static crystallographic references or docking scores alone, particularly in systems involving highly charged substrates and metal-dependent interactions.

Overall, these findings suggest that dynamically stabilized ligand accommodation within PPH-like cavities may provide a useful criterion for prioritizing antiparasitic scaffolds, particularly in systems involving flexible catalytic environments and metal-dependent substrate coordination. Although aptosimon emerged from the virtual screening stage, hinokinin was selected for experimental evaluation because it was available for biological testing and belonged to the broader prioritized natural-product chemical space. Accordingly, the cellular assays should be interpreted as scaffold-level biological validation rather than as direct validation of aptosimon as a single active compound.

Hinokinin has been described as a bioactive lignan with reported anti-inflammatory, antimicrobial, cytotoxic, and particularly antitrypanosomal properties [96]. Previous studies have also shown that free or formulated hinokinin can display activity against *T. cruzi*, including a hinokinin-loaded microparticle formulation that significantly reduced parasitemia in infected mice [92].

A limitation of the present study is that the cellular assays demonstrate antiparasitic activity of hinokinin but do not directly confirm biochemical inhibition of PPH. Accordingly, the proposed relationship between PPH ligand accommodation and cellular activity should be interpreted as a target-guided mechanistic hypothesis rather than as direct target validation. Future enzymatic inhibition assays, target-engagement studies, and, if feasible, genetic validation will be required to establish whether PPH is directly modulated by this scaffold in parasite cells.

## Conclusions

The present study combined proteome mining, phylogenetic analysis, structural modeling, molecular dynamics refinement, virtual screening, and experimental validation to investigate PPH as a potential therapeutic target in kinetoplastid parasites. Starting from a proteome-wide analysis of *Trypanosoma cruzi*, PPH emerged as a metabolically relevant and structurally tractable candidate, while comparative analyses demonstrated that related PPH-like proteins are conserved across kinetoplastids, including *L. donovani*. Because experimentally resolved structures were unavailable for the selected targets, AlphaFold-derived models were refined through extensive MD simulations, generating dynamically equilibrated conformations suitable for structure-based analyses. The refinement process showed that although the overall enolase-like fold remained conserved, important species-dependent differences existed in local flexibility, pocket accessibility, and catalytic-site organization. These differences became particularly evident during comparative analyses involving PPH and divalent metal coordination.

The integration of SiteMap analysis, docking, MM-GBSA calculations, and long-timescale MD simulations demonstrated that the selected binding environments could accommodate chemically diverse ligands beyond canonical substrate-like configurations. Importantly, ligand behavior observed during molecular dynamics simulations did not always correlate directly with static docking scores, emphasizing the importance of incorporating dynamic structural evaluation into structure-based antiparasitic screening workflows.

Among the evaluated compounds, the aptosimon-related lignan chemotype, represented experimentally by hinokinin, exhibited favorable interaction behavior and measurable antiparasitic activity. The compound demonstrated intracellular activity against *T. cruzi* amastigotes and maintained energetically favorable interaction profiles within both *T. cruzi* and *L. donovani* PPH-like models. These findings suggest that non-substrate-like natural-product-derived scaffolds may interact with conserved or functionally relevant regions of kinetoplastid PPH proteins without requiring strict reproduction of the endogenous catalytic configuration.

The comparative analyses also highlighted the limitations of directly extrapolating catalytic-site behavior across related kinetoplastid species. Although PPH conservation supported cross-species target exploration, local structural differences substantially influenced ligand accommodation and substrate-associated configurations. These observations reinforce the importance of species-specific structural refinement and dynamic evaluation when extending computational drug discovery pipelines across phylogenetically related parasites.

A limitation of the present study is that the cellular assays demonstrate antiparasitic activity of hinokinin but do not directly confirm biochemical inhibition of PPH. Therefore, hinokinin should be interpreted as an experimentally active natural product-derived scaffold emerging from the prioritized chemical space, rather than as a confirmed PPH inhibitor. Future enzymatic assays, target-engagement studies, or genetic validation approaches will be required to establish whether PPH is directly modulated by this scaffold in parasite cells.

Overall, this study establishes an integrated proteome-to-structure workflow for prioritizing PPH-like enzymes as candidate targets in kinetoplastid parasites. The results support further optimization of the aptosimon-related lignan chemotype represented by hinokinin and highlight the importance of combining omics-guided target selection, apo structural refinement, ligand-bound MD, and biological validation when evaluating natural-product-derived antiparasitic candidates.

## Author Contributions

Conceptualization, L.D.G.M., M.A.C.F.; methodology, L.D.G.M., H.L.B.C. M.N., L.P., C.S., and validation, L.D.G.M., H.L.B.C., M.A.C.F., L.P., M.N.; formal analysis, L.D.G.M., M.N., H.L.B.C., M.A.C.F.; investigation, L.D.G.M., H.L.B.C., M.N., and resources, C.S., M.A.C.F.; writing—original draft preparation, L.D.G.M., M.A.C.F.; writing—review and editing, L.D.G.M., H.L.B.C., M.N., L.P., M.A.C.F., C.S., and supervision, C.S., M.A.C.F. All authors have read and agreed to the published version of the manuscript.

## Funding

This work was funded by the Consejo Nacional de Ciencia, Tecnología e Innovación Tecnológica (CONCYTEC), the Programa Nacional de Investigación Científica y Estudios Avanzados (PROCIENCIA), by the call “E067-2023-01 Proyectos Especiales: Proyectos de Incorporación de Investigadores Postdoctorales en Instituciones Peruanas” [número de contrato PE501084367-2023], and Grant PE501088204-2024. Also, it was funded by the Universidad Católica de Santa María (grants 27499-R-2020, 27574-R-2020, 7309-CU-2020, and 28048-R-2021) and by the Research Management Office from the Universidad Católica de Santa María.

## Institutional Review Board Statement

Not applicable.

## Informed Consent Statement

Not applicable.

## Data Availability Statement

The software used in this study includes Maestro (Schrödinger Suite), Desmond, SiteMap, and Glide, all of which require a paid license for both academic and commercial use. Additionally, STRING, Cytoscape, and PASSer were utilized, which are freely available for non-commercial use. All software tools are referenced throughout the Methods and Results sections.

Furthermore, protein structures, docking grids, molecular dynamics trajectories, ligand preparation files, energy minimization data, and analysis scripts generated during this study are documented accordingly. All simulation scripts, docking results, molecular interaction analyses, and processed datasets can be found at https://github.com/CompBioChemRG.

## Acknowledgments

The authors express their gratitude for the financial support from the Programa Nacional de Investigación Científica y Estudios Avanzados—PROCIENCIA (PE501088204-2024 and PE501084367-2023) and the SNI Panama for partial funding to C.S.

## Conflicts of Interest

The authors declare no conflicts of interest.

## Supporting information

Supplementary material includes stereochemical validation of apo *T. cruzi* PPH frame selection (Supplementary Figure S1), FragFp-based structural similarity analysis of selected natural product candidates (Supplementary Figure S2), structural validation metrics for apo refinement frames (Supplementary Table S1), a summary of apo and ligand-bound MD systems (Supplementary Table S2), comparative hotspot persistence across *T. cruzi* ligand-bound simulations (Supplementary Table S3), and predicted ADME/drug-likeness descriptors for aptosimon and hinokinin (Supplementary Table S4).

## Notes

### Competing Interest Statement

The authors have declared no competing interest.

