## Supplementary material for "From Proteome Mining to Structural Validation: Phosphopyruvate Hydratase as a Structurally Tractable Drug Target in Kinetoplastid Parasites"

1. Computational Biology and Chemistry Research Group, Vicerrectorado de Investigación, Universidad Católica de Santa María, Arequipa 04000, Peru.

2. Center of Cellular and Molecular Biology of Diseases, Instituto de Investigaciones Científicas y Servicios de Alta Tecnología (INDICASAT AIP), City of Knowledge, Clayton, Apartado 0843-01103, Panama City, Panama.

3. DIFACQUIM Research Group, Department of Pharmacy, School of Chemistry, Universidad Nacional Autónoma de México, Mexico City, Mexico.

4. Instituto de Patobiología Veterinaria, CICVyA, Instituto Nacional de Tecnología Agropecuaria (INTA), Buenos Aires, Argentina.

5. Consejo Nacional de Investigaciones Científicas y Técnicas (CONICET), Buenos Aires, Argentina.

6. Programa de Pós-Graduação em Ciências da Saúde: Infectologia e Medicina Tropical, Faculdade de Medicina, Universidade Federal de Minas Gerais, Belo Horizonte, Brazil.

7. Departamento de Patologia Clínica, Colégio Técnico da Universidade Federal de Minas Gerais (COLTEC), Universidade Federal de Minas Gerais, Belo Horizonte, Brazil.

CONFIDENTIAL

**Abstract**

Chagas disease, caused by *Trypanosoma cruzi*, demands novel therapeutic strategies that overcome the toxicity and limited efficacy of current treatments. To address this need, herein we report an integrative, target-centric strategy that combines parasite proteome mining, structural modeling, and experimental validation. Functional enrichment and druggability analyses identified phosphopyruvate hydratase (PPH) as a promising candidate due to its essential metabolic role and limited similarity to human homologs. Notably, proteome mining revealed the presence and conservation of PPH across kinetoplastid parasites, including *Leishmania donovani*, supporting its evaluation beyond *T. cruzi*. For the selected PPH sequences, AlphaFold-derived three-dimensional models underwent extensive molecular dynamics refinement, yielding stable conformational ensembles suitable for structure-based studies. Using this validated model, virtual screening of the Latin American Natural Products Database - LANaPDB - identified aptosimon as a top-ranked compound candidate. Molecular dynamics simulations further showed ligand-dependent binding behavior, suggesting alternative binding modes distinct from the canonical substrate configuration. *In vitro* assays demonstrated consistent antiparasitic activity against intracellular *T. cruzi* amastigotes ($IC_{50}$ = 3.52 ± 0.023 µg/mL) and *Leishmania donovani* promastigotes ($IC_{50}$ = 13.06 ± 0.018 µg/mL), supporting the biological relevance of the aptosimon-related lignan chemotype, hinokinin, across two kinetoplastid parasite models. Together, these results support PPH as a structurally tractable and biologically relevant candidate target, while identifying an aptosimon-related lignan chemotype, represented experimentally by hinokinin, as a cross-species antiparasitic scaffold that warrants further biochemical target-validation studies.

**Keywords:** Chagas disease, Leishmaniasis, *Trypanosoma cruzi*, *Leishmania donovani*, Neglected tropical diseases, Structural modeling, Molecular dynamics, Natural product drug discovery, Aptosimon, Hinokinin, Phosphopyruvate hydratase.

**Comentado [CS1]:** Si estás de acuerdo, se podría mencionar esta otra clase de compuestos extraídos de las secreciones del torso de los sapos, no por citar un trabajo nuestro, sino por establecer la credibilidad de nuestro instituto en la búsqueda de compuestos activos de la naturaleza. De este tema hemos puboicado ya 3 papers, y vienen más en camino. Claro que tenemos muchísimos más dentro de las familias que ya mencionas, pero este, aunque se podría argumentar que de alguna manera es un terpenoide, es muy diferente. Si te parece bien, esta sería la mejor cita: Rodriguez C, Ibáñez R, Olmedo DA, Ng M, Spadafora C, Durant-Archibold AA, Gutiérrez M. Anti-Trypanosomal Bufadienolides from the Oocytes of the Toad *Rhinella alata* (Anura, Bufonidae). Molecules. 2023 Dec 29;29(1):196. doi: 10.3390/molecules29010196. PMID: 38202779; PMCID: PMC10779871

CONFIDENTIAL

$$\Delta G_{bind} = G_{complex} - (G_{protein} + G_{ligand})$$

where $G_{complex}$ corresponds to the free energy of the protein–ligand complex, and $G_{protein}$ and $G_{ligand}$ represent the energies of the isolated protein and ligand, respectively, calculated under identical conditions.

CONFIDENTIAL

with the crystallographic enolase from *Trypanosoma brucei* (PDB ID: 2PTY), where $Zn^{2+}$ ions were replaced with $Mg^{2+}$ ions to represent physiologically relevant cofactors. Ligand and metal positioning were defined before solvation based on the aligned structures.

All complexes were solvated in an explicit TIP3P water model within an orthorhombic simulation box, maintaining a minimum buffer distance of 10 Å between the protein and box boundaries. Systems were neutralized with counterions and adjusted to a physiological ionic strength of 0.15 M NaCl. The OPLS4 force field was applied to all systems.

Before production simulations, each system underwent a multistep relaxation protocol consisting of four steps, namely: (i) energy minimization, (ii) restrained NVT equilibration at 300 K for 1.0 ns with harmonic restraints applied to the protein backbone and ligand heavy atoms (10 $kcal\cdot mol^{-1}\cdot Å^{-2}$), (iii) NPT equilibration at 309.65 K and 1 atm for 1.0 ns with reduced restraints (2 $kcal\cdot mol^{-1}\cdot Å^{-2}$), and (iv) an additional 1.0 ns equilibration stage with minimal restraints (1 $kcal\cdot mol^{-1}\cdot Å^{-2}$).

Production MD simulations were then performed under NPT conditions (309.65 K, 1 atm) for 300 ns without positional restraints. Temperature was maintained using the Nose–Hoover thermostat, and pressure was controlled using the Martyna–Tobias–Klein barostat. Long-range electrostatics were treated with the particle mesh Ewald (PME) method, and a 2 fs time step was used with bond constraints applied to hydrogen atoms.

Trajectories were recorded at regular intervals for analysis. Structural stability was evaluated using root mean square deviation (RMSD) of protein backbone and ligand heavy atoms, while residue-level flexibility was assessed through root mean square fluctuation (RMSF). Protein–ligand interactions were monitored over time using interaction timeline analysis tools implemented in Schrödinger Maestro.

Identical simulation conditions were applied across all systems to ensure comparability between *T. cruzi* and *L. donovani* complexes.

A summary of apo-refinement and ligand-bound MD systems, simulation lengths, and representative frames is provided in Supplementary Table S2.

### Antiparasitic Activity Assays

#### Anti-*T. cruzi* intracellular assay

The anti-*Trypanosoma cruzi* activity was evaluated using the Tulahuen LacZ clone C4, which expresses β-galactosidase as a viability reporter [59]. Parasites were maintained in RPMI-1640 medium supplemented with L-glutamine, HEPES buffer, $NaHCO_3$, fetal bovine serum, and penicillin–streptomycin at 37 °C.

One day before infection, Vero cells were seeded in 96-well plates at $1.26 \times 10^4$ cells/well in a final volume of 100 µL of culture medium. After 24 h, cells were infected with $5 \times 10^4$ parasites/well diluted in 50 µL of culture medium. After an additional 24 h of infection, test compounds dissolved in DMSO were added and incubated for 120 h at 37 °C [60].

Antitrypanosomal activity was determined using chlorophenol red-β-D-galactopyranoside (CPRG; Roche Applied Science) as substrate for parasite β-galactosidase. CPRG was added to each well

CONFIDENTIAL

and allowed to react for 4.5 h at 37 °C. Absorbance was measured at 570 nm using a microplate reader [61]. Benznidazole was used as a positive control, and DMSO-treated infected cells were used as the negative control. $IC_{50}$ values were calculated from dose–response curves. In parallel, cytotoxicity was evaluated in non-infected Vero cells exposed to the same concentration range of hinokinin under comparable culture conditions. The $CC_{50}$ value was estimated as the concentration required to reduce Vero-cell viability by 50% relative to DMSO-treated controls and was used to calculate apparent selectivity indices as $SI=CC_{50}/IC_{50}$ [60].

**Anti-*Leishmania donovani* Promastigote Assay**

The anti-*Leishmania* activity was evaluated against *L. donovani* promastigotes following the protocol described by Calderón et al. [62], using the fluorescent DNA intercalator PicoGreen (Invitrogen, USA). This assay was used as an initial parasite-growth inhibition screen for the selected compound.

For each bioassay, 1 × $10^6$ promastigotes/well were placed in 96-well plates with test compounds diluted in DMSO, in a final volume of 100 µL, and incubated for 3 days. Amphotericin B was used as the positive control [63].

After incubation, a PicoGreen cocktail was added at a 1:4 dilution and incubated at room temperature for 5 min. Fluorescence was then measured at 485 nm. Antileishmanial activity was expressed as percent inhibition of growth relative to the DMSO-treated negative control. $IC_{50}$ values were calculated from serial dilution assays.

For hinokinin, serial dilutions of 50, 25, 12.5, 6.25, and 3.125 µg/mL were tested in the antiparasitic assays and in the Vero-cell cytotoxicity assay. $IC_{50}$ and $CC_{50}$ values were estimated from dose–response data and expressed as the concentrations required to reduce parasite growth or Vero-cell viability by 50%, respectively. Apparent selectivity indices were calculated as $SI = CC_{50}/IC_{50}$, using the Vero-cell $CC_{50}$ and the corresponding antiparasitic $IC_{50}$ values.

**Results and Discussion**

Figure 1 summarizes the complete methodological workflow, from proteome mining and target prioritization to structural refinement, ligand screening, MD simulations, and experimental evaluation of selected natural products.

CONFIDENTIAL

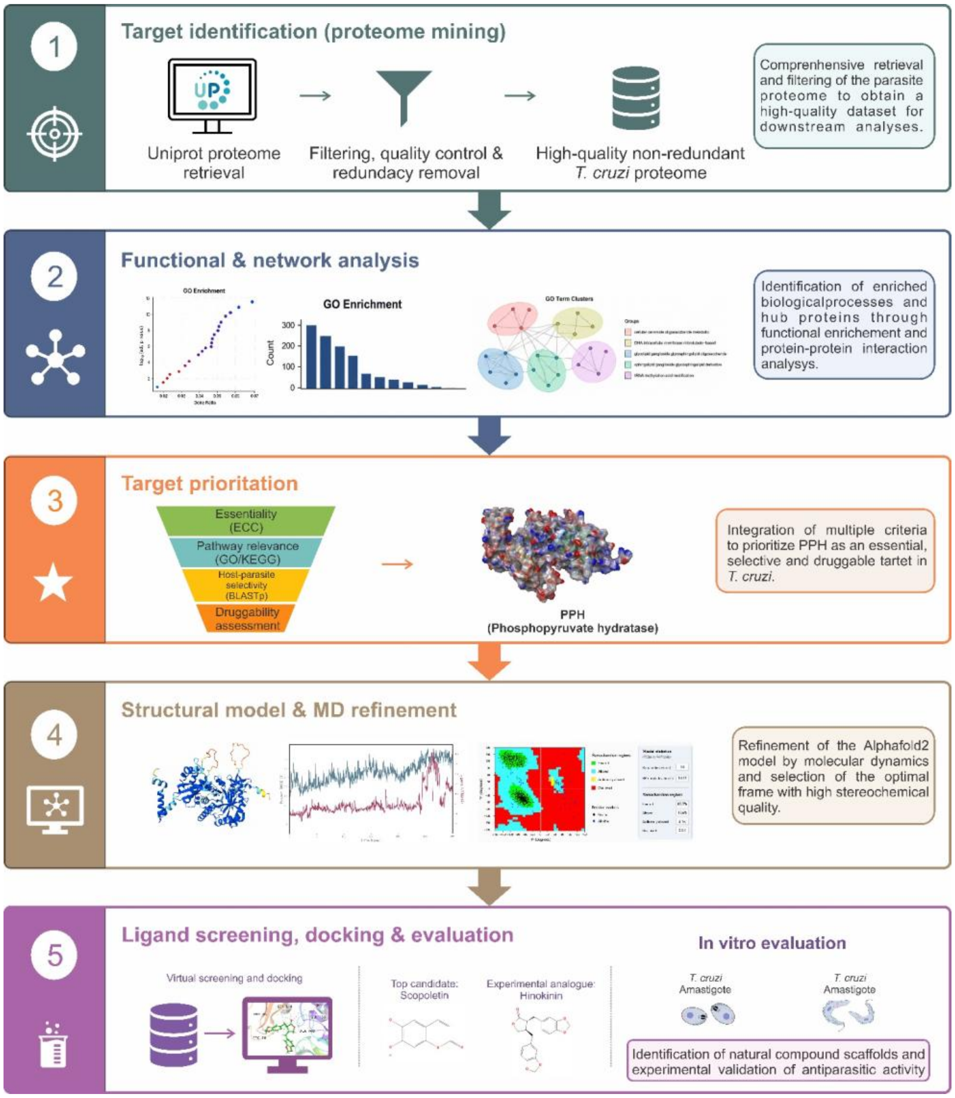


**Figure 1.** Integrated workflow for proteome-guided target prioritization, structural refinement, ligand screening, and experimental validation. The study integrated *T. cruzi* proteome mining, functional and network analysis, PPH prioritization, AlphaFold-based structural modeling, apo MD refinement, binding-site identification, virtual screening, ligand-bound MD simulations, and *in vitro* testing using *T. cruzi* and *Leishmania donovani* assays.

### Proteome Mining, Functional Enrichment, and Target Prioritization

The proteome-wide analysis of *T. cruzi* identified a functionally diverse set of biological processes potentially relevant for parasite survival, adaptation, and therapeutic intervention. Functional enrichment analysis highlighted biological process categories associated with RNA modification, lipid metabolism, carbohydrate metabolism, intracellular organization, microtubule-associated processes, intracellular signaling, metabolic regulation, and overlapping functional categories associated with parasite adaptation and viability (Figure 2A-C). These enriched categories collectively indicate that *T. cruzi* depends on tightly coordinated metabolic and regulatory systems that support translational fidelity, membrane remodeling, intracellular adaptation, and energetic plasticity throughout its complex life cycle [64]. Rather than representing isolated pathways, the enrichment profile suggests the existence of highly interconnected biological networks required for parasite survival across distinct environmental and host-associated conditions. This system-level organization is notable from a therapeutic perspective because disruption of central metabolic hubs may propagate downstream effects across multiple adaptive pathways simultaneously.

The bar and upset plots further demonstrated that several enriched biological processes were partially interconnected rather than functionally isolated, supporting the existence of coordinated metabolic and regulatory modules associated with parasite survival and adaptation (Figure 2B).

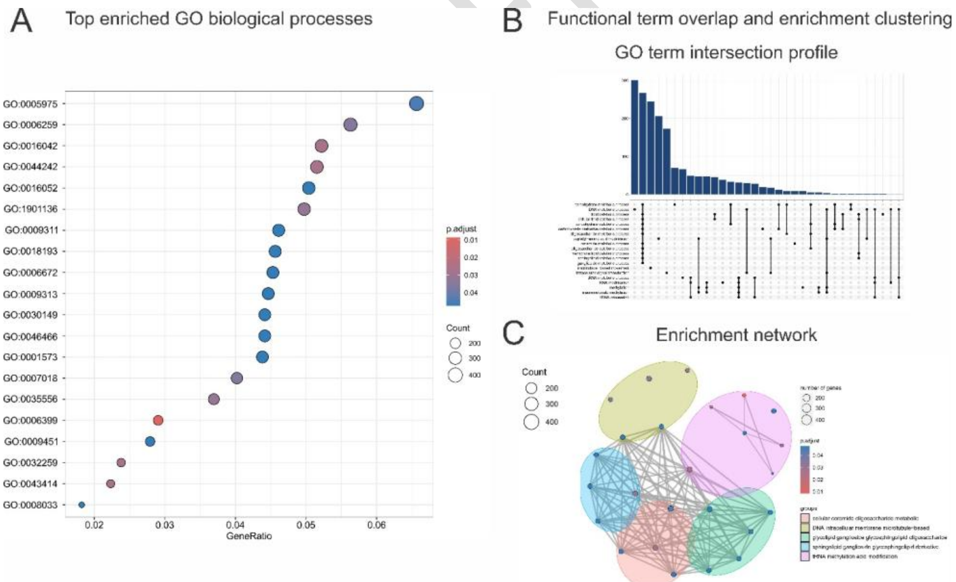


**Figure 2.** Functional enrichment analysis of *T. cruzi* proteins. (A) Dot plot highlights top Gene Ontology terms by gene ratio and adjusted p-value. (B) The bar and upset plots (top) show the distribution and overlap of enriched biological processes. (C) The network plot (bottom) groups

related pathways emphasize key clusters in tRNA modification, lipid metabolism, and microtubule organization.

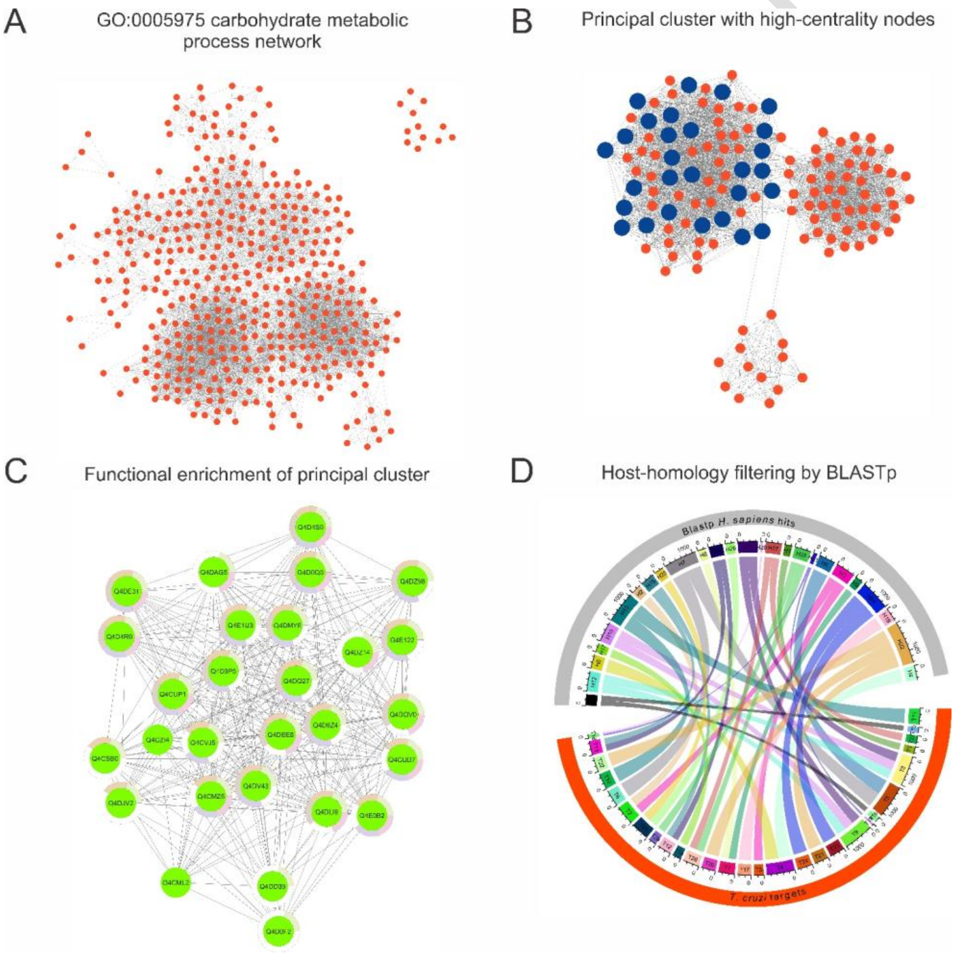

**Figure 3. Refined network analysis of the carbohydrate metabolic process GO:0005975.** (A) Protein–protein interaction network associated with GO:0005975, selected from the enrichment analysis shown in Figure 2. (B) Principal interaction cluster extracted from this network, with high-priority/high-centrality nodes highlighted in orange. (C) Functional enrichment analysis of the highlighted nodes, showing pathway categories linked to central metabolism, including glycolysis/gluconeogenesis, carbon metabolism, and the pentose phosphate pathway. (D) BLASTp-based comparison of prioritized parasite proteins against *Homo sapiens* homologs, supporting host-similarity filtering and prioritization of candidate targets, including phosphopyruvate hydratase (PPH) and phosphoglucomutase (PGM).

Together, the GO:0005975 network refinement, cluster enrichment, and BLASTp-based host-homology filtering supported the prioritization of metabolically relevant parasite candidates with potential therapeutic selectivity.

Within this framework, phosphopyruvate hydratase (PPH) (UniProt ID: Q4DMY6), also known as enolase, emerged as a prioritized candidate. PPH catalyzes the reversible conversion of 2-phosphoglycerate into phosphoenolpyruvate, a key step in glycolysis [78], placing it within an essential energy-producing pathway. Beyond its canonical metabolic function, enolase has also been associated with non-glycolytic or "moonlighting" roles in several pathogens, including participation in host–parasite interactions and surface-associated functions [79]. This multifunctional behavior increases the biological relevance of PPH, as its inhibition could potentially affect parasite energy metabolism and additional parasite-specific adaptive processes linked to infection [80–82]. This observation provided the rationale for extending the downstream structural and ligand-binding analyses to a *L. donovani* PPH-like model as a comparative kinetoplastid system. This extension was not based on epidemiological overlap, but on the conservation of PPH/enolase-like enzymes across trypanosomatid parasites and the opportunity to evaluate whether ligand-accommodation features identified in *T. cruzi* could retain relevance in a related parasite model. The phylogenetic distribution observed across kinetoplastids therefore supported a comparative cross-species analysis of PPH-like structural features and ligand-binding behavior (Figure 4A) [83] [84]. The biological, structural, and selection criteria supporting PPH prioritization, together with the rationale for including the comparative *L. donovani* PPH-like model, are summarized in Table 1.

CONFIDENTIAL

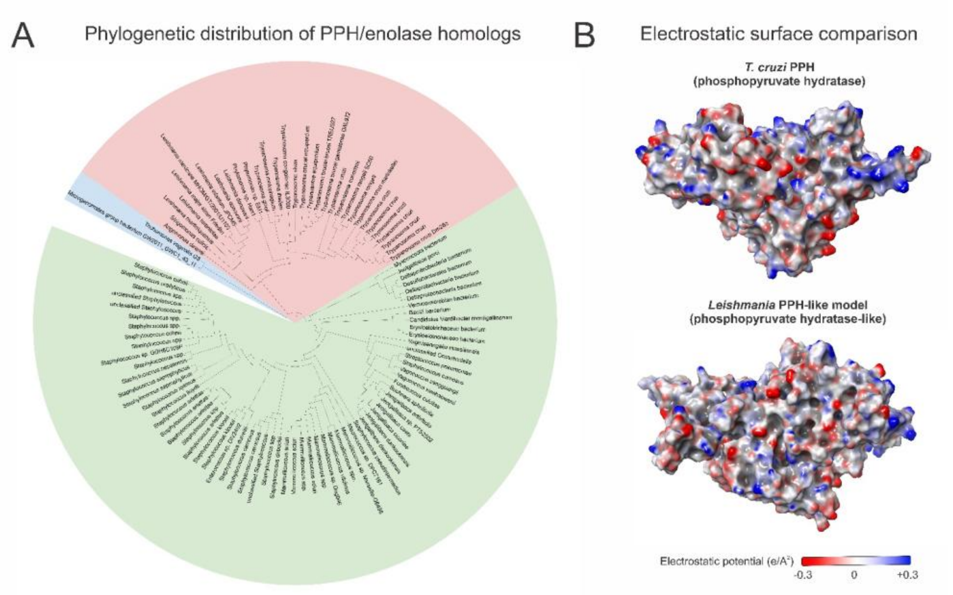


**Figure 4.** Phylogenetic distribution and electrostatic surface features of PPH-like enzymes. (A) Phylogenetic tree showing conservation of phosphopyruvate hydratase/enolase homologs across kinetoplastid parasites and broader eukaryotic taxa. Green sectors denote kinetoplastid species, whereas red sectors represent non-kinetoplastid eukaryotic groups. (B) Electrostatic surface potential maps of the MD-refined *T. cruzi* PPH model and the *Leishmania* PPH-like model, highlighting species-dependent charge distribution around surface-accessible regions used for downstream pocket and ligand-accommodation analyses.

**Table 1.** Rationale for PPH prioritization and comparative structural evaluation**.** Summary of the biological, structural, and selection criteria supporting the prioritization of *T. cruzi* PPH and the inclusion of a *L. donovani* PPH-like model for comparative ligand-accommodation analyses.

| Target | Species | Biological role | Structural rationale | Selection rationale |
|---|---|---|---|---|
| PPH (Phosphopyruvate hydratase) | *T. cruzi* | Central glycolytic enzyme associated with parasite energy metabolism | Conserved catalytic architecture with identifiable ligand-accommodating hotspot | Selected for comparative structural and dynamic investigation |
| PPH-like model | *L. donovani* | Predicted glycolytic/metabolic function | Partial catalytic transferability with preserved adaptable ligand region | Included as a comparative kinetoplastid model to evaluate conservation of PPH-like ligand- |

CONFIDENTIAL

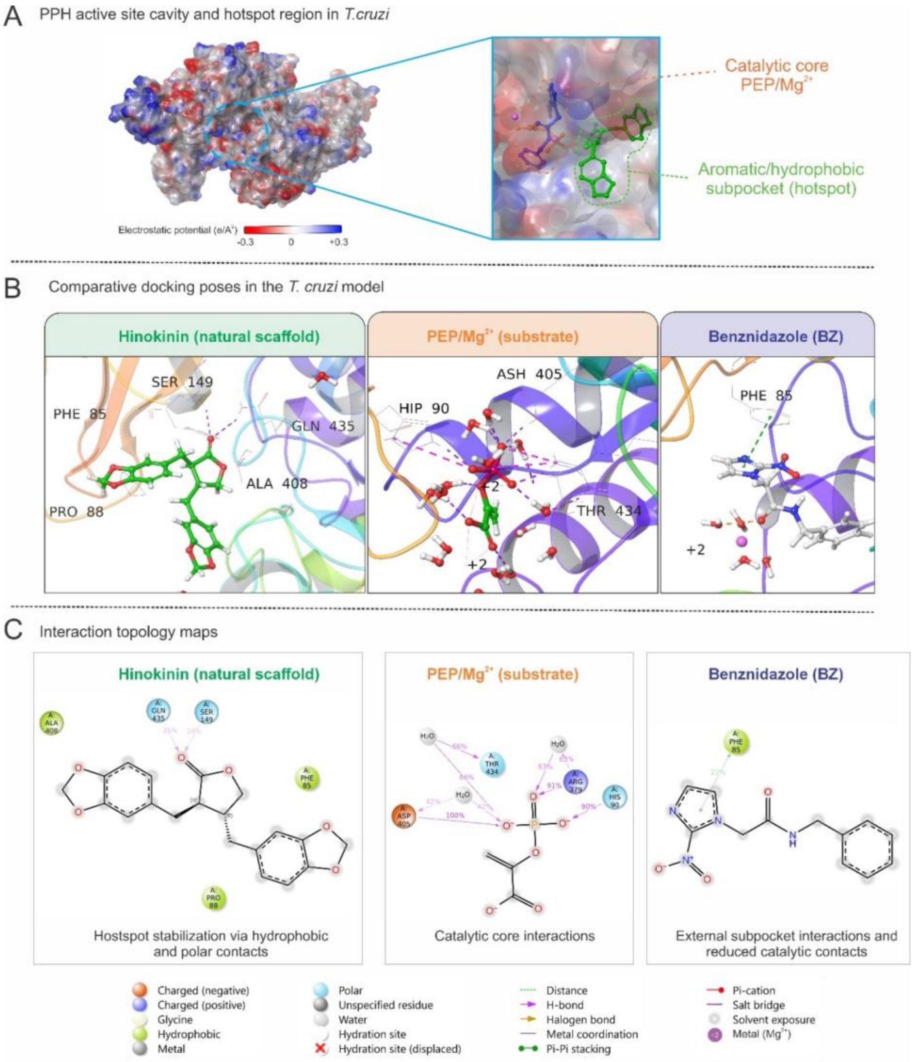


**Figure 6**. Binding-site characterization and ligand-accommodation behavior in *T. cruzi* PPH. (A) SiteMap-defined pocket and catalytic-reference mapping using the *T. brucei* enolase PEP/$Mg^{2+}$ complex. (B) Comparative docking poses of PEP/$Mg^{2+}$, hinokinin (aptosimon-like lignan analogue), and benznidazole within the selected cavity. (C) Two-dimensional interaction diagrams showing ligand-specific contact with the catalytic core and adjacent hotspot regions. Aptosimon was prioritized during virtual screening, whereas hinokinin was used as an experimentally accessible, structurally related scaffold for biological evaluation.

Structural superposition with the crystallographic *T. brucei* enolase complex revealed that the selected cavity partially overlapped with the substrate-associated region occupied by phosphoenolpyruvate (PEP) and divalent metal ions in the reference structure [48]. This overlap reinforced the functional relevance of the selected binding site and supported its use for docking and molecular dynamics analyses. Replacement of crystallographic $Zn^{2+}$ ions with $Mg^{2+}$ ions allowed construction of catalytic-like configurations compatible with the known metal dependence of enolase enzymes [83]. Visually, the resulting structural arrangement suggested the coexistence of a polar catalytic region associated with PEP/$Mg^{2+}$ coordination and an adjacent mixed hydrophobic–polar cavity capable of accommodating non-substrate-like ligands (Figure 6A–C).

Principal component analysis (PCA) of the screened compounds revealed a broad and heterogeneous distribution of chemical space across the different LANaPDB-associated sources, including $\text{NuBBE}_{\text{DB}}$, BIOFACQUIM, SistematX, UEFS, NAPRORE, UNIIQUIM, NPDB EjeCol, CIFPMA, PeruNPDB, and LAIPNUDELSAV (Figure 5A). The observed dispersion indicates that the dataset encompasses structurally diverse natural products rather than highly redundant chemical families, an important feature for exploratory drug discovery. Compounds derived from databases occupied partially overlapping but distinguishable regions of the PCA space, suggesting the coexistence of both shared and unique molecular scaffolds among regional biodiversity collections. This chemical heterogeneity is particularly advantageous in exploration and antiparasitic discovery because structurally diverse scaffolds may access distinct interaction regions within flexible enzymatic cavities.

CONFIDENTIAL

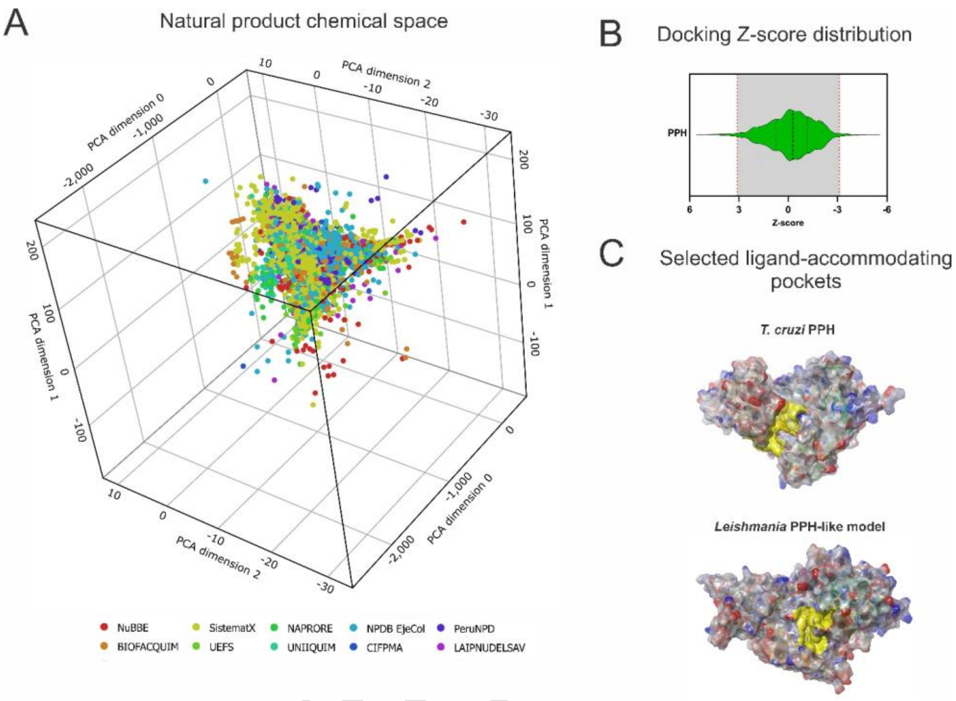


**Figure 5.** Chemical space distribution, docking-score prioritization, and pocket-centered screening strategy for PPH-like targets. (A) PCA-based chemical space distribution of the initial Latin American Natural Product Database included in the screening workflow. Colors indicate compound database source. (B) Docking Z-score distribution for compounds screened against PPH, highlighting the threshold-based prioritization region. (C) Electrostatic surface views of the MD-refined *T. cruzi* PPH and *L. donovani* PPH-like models, with SiteMap-selected ligand-accommodating pockets shown in yellow.

CONFIDENTIAL

superposition and replacement of $Zn^{2+}$ by $Mg^{2+}$, the modeled *T. cruzi* PEP–$Mg^{2+}$ system reproduced the general requirement for a polar, metal-dependent substrate-recognition environment, although the exact residue numbering and local geometry differed between species.

MD simulations further identified distinct binding profiles between the evaluated ligands. Benznidazole initially maintained interactions near the catalytic region but progressively migrated toward a more external region of the cavity during later stages of the trajectory. In contrast, the hinokinin-associated lignan analogue displayed comparatively stable residence within the selected binding region over the simulation timeframe. These differences reinforce the notion that docking scores alone may not fully capture the dynamic stability of ligand binding within the PPH cavity.

RMSD analysis indicated that the hinokinin-associated system stabilized rapidly within the first 10–15 ns and fluctuated predominantly within a ~0.35–0.60 nm range throughout the simulation, whereas the apo structure stabilized at higher average deviations (~0.55–0.65 nm). In contrast, the PEP/$Mg^{2+}$ system exhibited broader fluctuations and progressive destabilization of the catalytic arrangement during later simulation stages.

Residue-wise RMSF analysis revealed damped fluctuations around pocket-lining regions associated with ligand accommodation, including residues surrounding the aromatic stabilization region previously associated with Trp398-like interactions. These reduced fluctuations support persistent local stabilization of the binding environment throughout the trajectory.

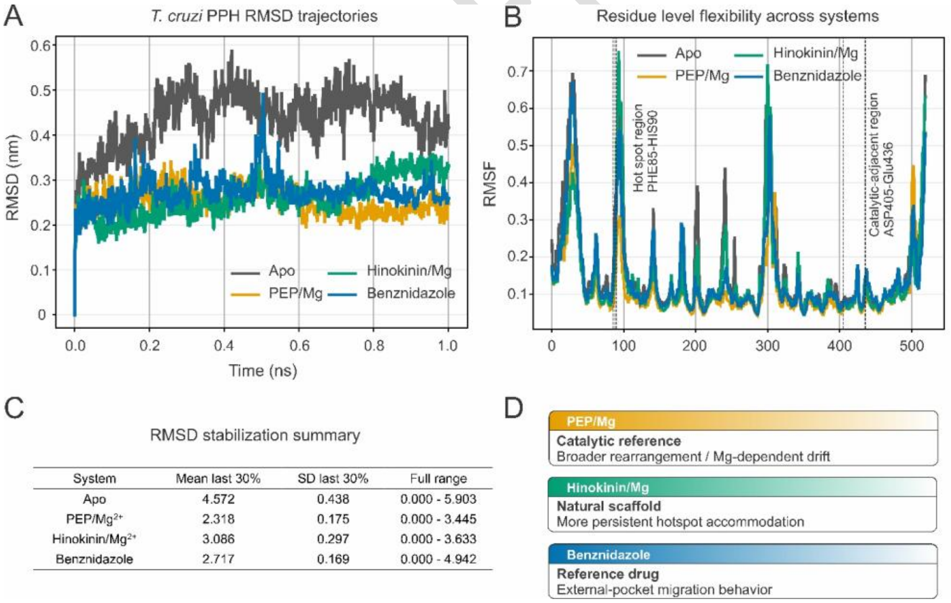


**Figure 7.** Ligand-bound molecular dynamic behavior of *T. cruzi* PPH complexes. (A) Backbone RMSD profiles of apo PPH and ligand-bound systems over 300 ns simulations, showing rapid stabilization of the hinokinin-associated complex and broader fluctuations in the PEP/$Mg^{2+}$

CONFIDENTIAL

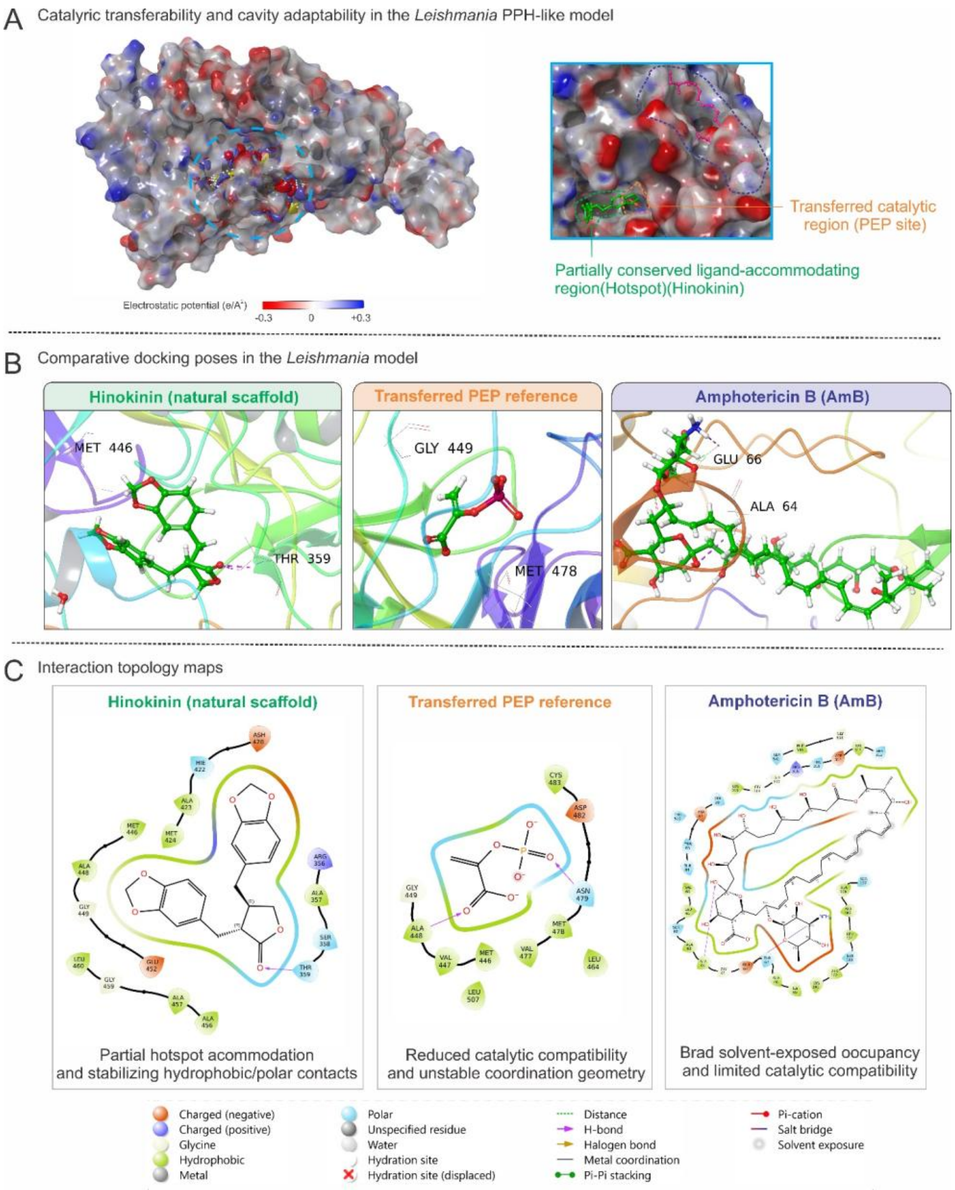


**Figure 8.** Comparative structural and ligand-accommodation behavior of the Leishmania PPH-like model. (A) MD-derived structural behavior of the Leishmania PPH-like model, highlighting increased flexibility in loop-rich and surface-accessible regions. (B) Structural mapping of the PEP/$Mg^{2+}$ catalytic reference from T. brucei enolase onto the Leishmania model, showing limited

transferability and partial burial of the substrate-like configuration. (C) Docking and interaction analyses showing favorable accommodation of hinokinin within a SiteMap-defined accessible cavity.

**Table 2.** Antiparasitic activity and apparent selectivity of hinokinin. $IC_{50}$ values are shown for hinokinin against intracellular *T. cruzi* amastigotes and *L. donovani* promastigotes, together with the $CC_{50}$ value determined in non-infected Vero cells. Apparent selectivity indices were calculated as SI = $CC_{50}/IC_{50}$ using the Vero-cell $CC_{50}$ and the corresponding antiparasitic $IC_{50}$ values.

| Compound | Parasite / assay | $IC_{50}$ (µg/mL) | Vero cell $CC_{50}$ (µg/mL) | Apparent SI |
|---|---|---|---|---|
| Hinokinin | *T. cruzi* intracellular amastigotes | 3.52 ± 0.023 | 36.89 ± 0.751 | ≈9.9 |
| | *L. donovani* promastigote | 13.06 ± 0.018 | 36.89 ± 0.751 | ≈2.8 |

Reference controls from the same dataset: benznidazole $IC_{50}$ = 0.81 µg/mL; amphotericin B $IC_{50}$ = 140.47 ng/mL.

17. Rodriguez, C.; Ibáñez, R.; Olmedo, D.A.; Ng, M.; Spadafora, C.; Durant-Archibold, A.A.;


CONFIDENTIAL

Gutiérrez, M. Anti-Trypanosomal Bufadienolides from the Oocytes of the Toad Rhinella Alata (Anura, Bufonidae). *Molecules* **2023**, *29*, 196.

24. Carlson, M.; Pagès, H. AnnotationForge: Tools for Building SQLite-Based Annotation Data Packages. R Package Version 1.42. 2 2023.

33. Gu, Z.; Gu, L.; Eils, R.; Schlesner, M.; Brors, B. " Circlize" Implements and Enhances

CONFIDENTIAL

Circular Visualization in R. **2014**.

CONFIDENTIAL

*274*, 5077–5089.

62. Calderon, A.I.; Romero, L.I.; Ortega-Barría, E.; Brun, R.; Correa A, M.D.; Gupta, M.P. Evaluation of Larvicidal and in Vitro. Antiparasitic Activities of Plants in a Biodiversity Plot

CONFIDENTIAL

in the Altos de Campana National Park, Panama. *Pharm. Biol.* **2006**, *44*, 487–498.

CONFIDENTIAL

249, doi:https://doi.org/10.1016/j.mib.2021.07.017.

CONFIDENTIAL


CONFIDENTIAL
